# Multi-omic rejuvenation of naturally aged tissues by a single cycle of transient reprogramming

**DOI:** 10.1101/2022.01.20.477063

**Authors:** Dafni Chondronasiou, Diljeet Gill, Lluc Mosteiro, Rocio G. Urdinguio, Antonio Berenguer, Monica Aguilera, Sylvere Durand, Fanny Aprahamian, Nitharsshini Nirmalathasan, Maria Abad, Daniel E. Martin-Herranz, Camille Stephan Otto-Attolini, Neus Prats, Guido Kroemer, Mario F. Fraga, Wolf Reik, Manuel Serrano

## Abstract

The expression of the pluripotency factors OCT4, SOX2, KLF4 and MYC (OSKM) can convert somatic differentiated cells into pluripotent stem cells in a process known as reprogramming. Notably, cycles of brief OSKM expression do not change cell identity but can reverse markers of aging in cells and extend longevity in progeroid mice. However, little is known about the mechanisms involved. Here, we have studied changes in the DNA methylome, transcriptome and metabolome in naturally aged mice subject to a single period of transient OSKM expression. We found that this is sufficient to reverse DNA methylation changes that occur upon aging in the pancreas, liver, spleen and blood. Similarly, we observed reversion of transcriptional changes, especially regarding biological processes known to change during aging. Finally, some serum metabolites altered with aging were also restored to young levels upon transient reprogramming. These observations indicate that a single period of OSKM expression can drive epigenetic, transcriptomic and metabolomic changes towards a younger configuration in multiple tissues and in the serum.

## MAIN TEXT

The simultaneous expression of four specific factors, OCT4, SOX2, KLF4 and MYC (OSKM), also known as “Yamanaka factors”, in adult differentiated cells is able to shut off their transcriptional programs for cell identity, activate the transcription of pluripotency genes, and establish a new identity equivalent to embryonic stem cells^1^. This process, generally known as reprogramming, erases molecular and cellular traits of aging acquired by somatic cells throughout their lifespan^2–6^. Moreover, reprogrammed cells can subsequently differentiate into somatic cells that are now rejuvenated relative to their parental cells^2, 3^. Among the various molecular changes associated to aging, DNA methylation at specific CpG sites has turned out to be tightly linked to aging and, particularly, to biological aging rather than chronological aging^7, 8^. Interestingly, examination of aging-associated DNA methylation has revealed that rejuvenation of this aging trait occurs progressively during reprogramming, being initiated at the early stages of the process and continuing until full reprogramming^9, 10^.

The expression of OSKM in mice recapitulates the process of reprogramming^11–13^. Upon OSKM expression *in vivo*, a fraction of cells within tissues shut off their cell identity markers and progressively activate the pluripotency program^11^. The completion of reprogramming *in vivo* manifest by the formation of teratomas, a tumor overgrowth formed by pluripotent cells differentiating into multiple cell lineages^11, 12^. Of note, interruption of the process of reprogramming at its early stages is fully reversible and does not result in a detectable risk of teratoma. Even more remarkably, cycles of short OSKM expression followed by recovery result in rejuvenation, both *in vivo* and *in vitro*^14, 15^. Rejuvenation by multiple cycles of OSKM has been demonstrated at various levels. In cells, there is a reduction in aging-associated DNA damage and epigenetic alterations, including DNA methylation at specific CpG sites^14, 15^. In mice, cycles of OSKM in adult mice improve their capacity to respond to tissue injury^14^. Finally, cycles of OSKM in progeric mice with constitutive DNA damage extend significantly their lifespan^14^. More recently, viral transduction of OSK in the retina of old mice has been shown to reduce aging-associated DNA methylation and to improve vision^16^.

Here, we perform a multi-omic and multi-tissue analysis of naturally aged mice exposed to a single cycle of transient OSKM expression. By studying the effects of a single cycle of OSKM, we aim to capture the more direct effects of transient reprogramming. We also examine, not only DNA methylation changes associated to aging, but also transcriptomic and serum metabolites. Finally, we observe *in vivo* rejuvenation in the pancreas where OSKM is highly expressed, but, interestingly, also in liver, spleen and peripheral blood where OSKM is weakly expressed.

### OSKM promotes epigenetic rejuvenation in pancreas

To induce transient *in vivo* reprogramming, we used a previously reported strain of mice in which the expression of OCT4, SOX2, KLF4 and MYC (OSKM) can be temporarily induced by doxycycline supplementation in the drinking water^11, 13^. In these mice, the pancreas is the most susceptible tissue to undergo reprogramming, followed by the intestine and stomach^11^. Reprogrammed tissues present focal areas of dysplasia in which the tissue alters its normal architecture and cells lose differentiation markers^11^. At a more advanced stage, tissues present compact areas of undifferentiated cells expressing pluripotency markers, such as NANOG^11^. To evaluate the effects of OSKM activation in old mice, we treated 55 weeks old reprogrammable mice for one week with a low dose of doxycycline (0.2 mg/ml). The age of 55 weeks was chosen in an effort to detect age-related differences caused by functional decline, rather than to alterations in cell composition that often occur at older ages; and the specific conditions for the treatment with doxycycline were chosen to avoid the formation of teratomas according to our previous experience^11^. Upon OSKM induction for 1 week, we observed clear histological changes in the pancreas (**Extended Data Fig. 1a**). This treatment, however, was not sufficient to achieve pluripotency, as judged by minimal or undetectable expression of pluripotency markers *Nanog*, *Oct4* (endogenous), and *Tfe3* in pancreas (**Extended Data Fig. 1b,c**). Of note, the detection of pluripotency markers in pancreas required two weeks of OSKM expression (**Extended Data Fig. 1b**). Importantly, the histological changes observed in the pancreas after one week of OSKM induction were reversed two weeks after removal of doxycycline (**Extended Data Fig. 1a**). Therefore, one week of OSKM expression allows for transient and reversible changes in the histology of the pancreas without achieving pluripotency. It is important to mention that while pancreas, intestine and stomach manifest histological changes during OSKM expression, other tissues, like liver or spleen, do not present observable histological changes (**Extended Data Fig. 1a**). It is also worth mentioning that the OSKM cassette is not homogeneously expressed in all cell types. For example, acinar cells of the pancreas rapidly and broadly express SOX2, 24h after i.p. injection of doxycycline, as detected by immunohistochemistry; however, this was not the case of other cell types of the pancreas (**Extended Data Fig. 1d**).

Changes in the epigenome, and particularly DNA methylation, are robustly linked to aging^7, 8, 17, 18^. To address the impact of transient OSKM expression on epigenetic aging *in vivo*, we used *MspI*-based Reduced Representation Bisulfite Sequencing (RRBS)^19^. We performed this analysis on genomic DNA from pancreata of reprogrammable mice at the age of 55 weeks that had been treated as described above, referred to as “old-OSKM” group (n=5). As control groups (n=5 per group), we used reprogrammable mice of 55 weeks of age (“old” group) and of 13 weeks (“young” group) without treatment with doxycycline (**Extended Data Fig. 1e**). Of all the methylation sites probed by RRBS, we focused on those located at regulatory elements, particularly promoters and enhancers. In total, we identified 11.272 promoters, which represent about 35% of the active promoters in pancreas^20, 21^, defined as H3K27ac-rich regions around transcription start sites; similarly, we identified 5.737 enhancers, defined as non-promoter H3K27ac-rich regions^20, 21^, which represent about 42% of the active enhancers in pancreas. By comparing old and young groups, we identified a list of differentially methylated (DM) promoters (**Extended Data Fig. 1f**). Focusing on these aging-DM promoters, we performed Principal Component Analyses (PCA) that clearly segregated young from old mice. Interestingly, the methylation profile of old-OSKM mice was placed by PCA between the young and old groups, revealing partial epigenetic rejuvenation of aging-sensitive promoters (**Fig. 1a, Supplementary Information**). Furthermore, we divided these aging-DM promoters into two groups depending on their age-associated gain or loss of methylation, respectively. This yielded a subset of promoters that are hypermethylated with aging and demethylated by OSKM (19 out of 51 in total, 37%), and a subset of promoters that are hypomethylated with aging and remethylated by OSKM (17 out of 42 in total, 40%) (**Fig. 1b**, **Extended Data Fig. 1g-h, Supplementary Table 1**). As a notable example, the *Hnf1a* promoter, a key transcription factor for the development of the pancreas^22^, was among the promoters hypermethylated with aging and reset by OSKM (**Fig. 1b**).

**Figure 1.**
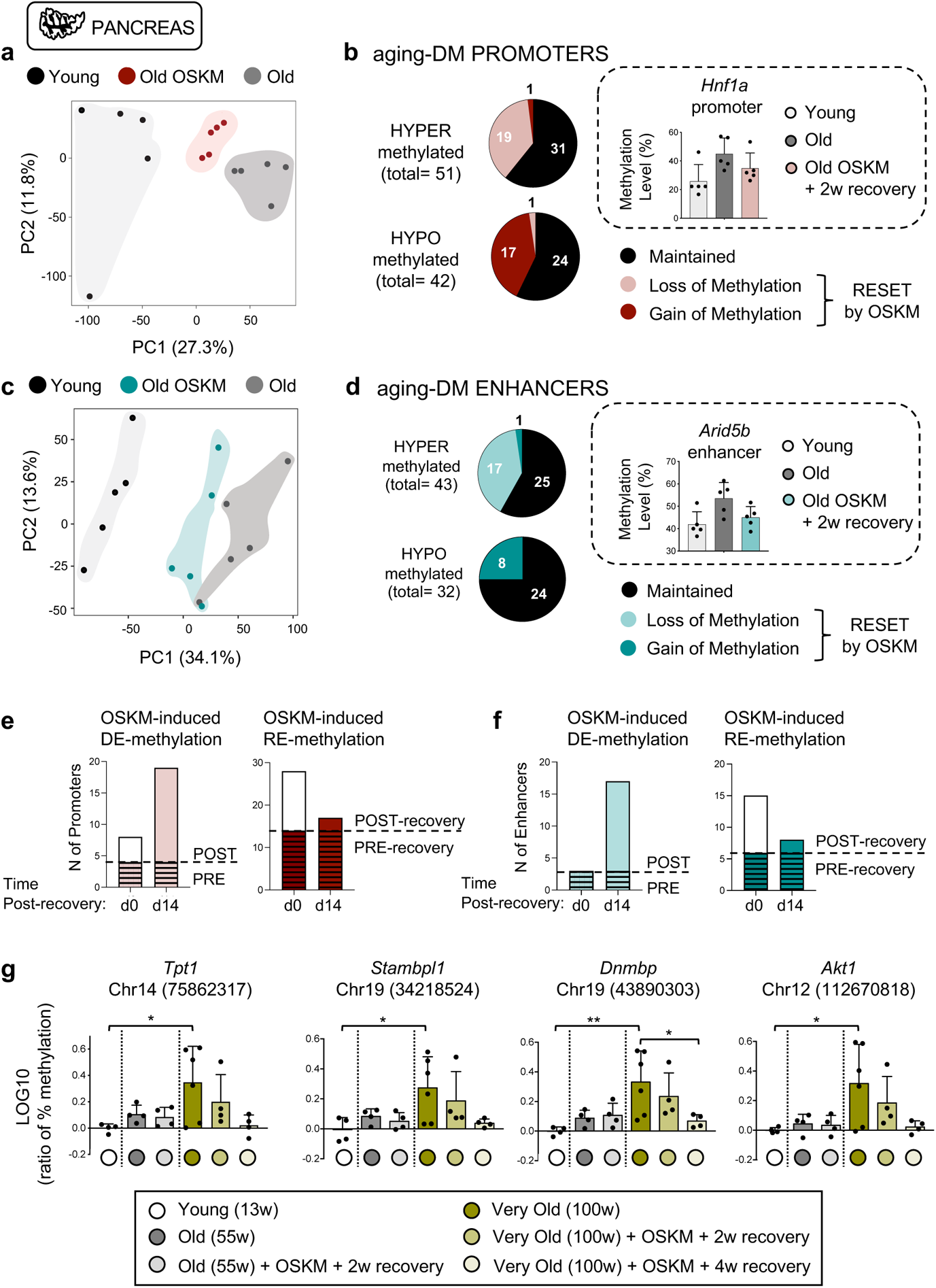
Transient OSKM reprogramming partially rejuvenates the methylation profile of old pancreas. **a,** Principal Component Analysis (PCA) of aging-related differentially methylated (DM) promoters of young (13 weeks, n=5), old (55 weeks, n=5) and old-OSKM (55 weeks, n=5) pancreas. **b,** DM promoters were divided into hyper- and hypo-methylated during aging, and shown is the number of these promoters that alters their methylation profile due to transient OSKM activation. *Hnf1a* promoter is a representative example of an age-associated hyper-methylated promoter that becomes demethylated in old-OSKM pancreas. **c,** PCA of aging-associated DM enhancers of young, old and old-OSKM pancreas. **d,** DM enhancers were classified into hyper- and hypo-methylated during aging, and shown is the subset of these enhancers that alters their methylation profile due to transient OSKM activation. *Arid5b* enhancer is a representative example of an age-related hyper-methylated enhancers that becomes demethylated in old-OSKM pancreas. **e,** The methylation status of aging-hypermethylated or hypomethylated promoters and **f,** enhancers that were found above to be OSKM-demethylated or remethylated respectively was evaluated directly after OSKM cessation (day 0 post-recovery) and 14 days post-recovery. **g,** Methylation levels measured by bisulfite pyrosequencing of four CpGs, located in regions hypermethylated with aging in the pancreas (n = 4 to 6). Bars in b, d and g represent the standard deviation (SD) of the data. Statistical significance was evaluated using one-way ANOVA with Tukey’s multiple comparison method, and comparisons are indicated as *P < 0.05 and **P < 0.01.

Similarly, we generated a list of differentially methylated enhancers by comparing the methylation levels of young *versus* old pancreata (**Extended Data Fig. 1i**). As before, PCA analysis showed that aging-DM enhancers in OSKM mice were distinct from old mice and displaced towards the young group (**Fig. 1c Supplementary Information**). We then separated those enhancers that are hypermethylated with aging and demethylated by OSKM (17 out of 43 in total), and those enhancers hypomethylated with aging and remethylated by OSKM (8 out of 32 in total) (**Fig. 1d, Extended Data Fig. 1j-k, Supplementary Table 2**). Of note, *Arid5a*, a pro-inflammatory RNA binding protein induced by NF-κB^23^, was identified as the gene in proximity to a DM enhancer (Chr10: 68231400-68232600) that was hypermethylated with aging but reduces its methylation after OSKM activation (**Fig. 1d**).

We wondered to what extent the observed changes in methylation occurred during the period of OSKM expression (1 week) or during the post-recovery period (2 weeks). For this, we analyzed the aging-DM regions in a group of 55 weeks old reprogrammable mice at the end of OSKM expression (d0 post-recovery) and compared it with the same regions after recovery (d14 post-recovery). We observed that the majority of OSKM-induced demethylation in the pancreas happened after turning off OSKM expression. In contrast, remethylation events were already present at d0 and half of them were preserved during recovery, while the other half were lost (**Fig. 1e,f**).

To confirm the effects of OSKM on methylation with a different technique, individual CpG sites were selected among the above-identified DM regions for validation by bisulfite pyrosequencing. For this validation, we used very old mice (around 100 weeks): wild-type mice treated with doxycycline (“very old”, n=6), and reprogrammable mice treated with doxycycline for one week followed by two weeks of recovery (“very old-OSKM+2w”, n=4) or four weeks of recovery (“very old-OSKM+4w”, n=4) (**Extended Data Fig. 1e**). We selected those individual CpGs within the RRBS with the highest methylation changes with aging and optimal sequence context for pyrosequencing (**Supplementary Table 3**). We tested a total of eleven CpGs. Among them, nine CpGs expected to gain methylation with aging (**Fig. 1g, Extended Data Fig. 1l,m**). Out of them, eight showed the expected increase in methylation with aging, specifically, those near genes *Tpt1* (2 close CpGs), *Stambpl1*, *Dnmbp* (2 consecutive CpGs), *Akt1*, *Gm12339* and *Gm17678*. Interestingly, all eight regions presented reduced methylation in very old-OSKM mice treated with doxycycline, in some cases becoming indistinguishable from young mice (**Fig. 1g, Extended Data Fig. 1l**). Of note, reversion of methylation was consistently more profound after 4 weeks of recovery than after 2 weeks of recovery. This, together with the RRBS data above (**Fig. 1e,f**), is another indication that the recovery period is important for the demethylation of aging-DMRs. We also tested two aging-hypomethylated CpGs (*1700011F14Rik* and *Ptprj*) and pyrosequencing confirmed that both were hypomethylated with aging. However, we could not detect remethylation in these two positions after transient OSKM activation (**Extended Data Fig. 1n**).

In summary, RRBS analysis has revealed a total of 93 promoters and 75 enhancers in which methylation changes with aging, and out of them a total of 61 (36%) were reversed towards a younger state by OSKM expression in old mice (**Fig. 1b,d**). Using a separate cohort of old mice and a different technique (pyrosequencing), we confirmed rejuvenation in 8 out of 11 tested regions that showed aging-dependent methylation changes. We conclude that a single period of OSKM expression is able to rejuvenate a fraction of the methylation changes that occur with aging in promoters and enhancers.

### Transcriptional rejuvenation of the pancreas

To analyze the effects of aging and OSKM at the transcriptional level, we performed RNAseq of the pancreata of young (13 weeks, n=4), old (55 weeks, n=5) and old-OSKM (55 weeks, 1 cycle of OSKM followed by 2 weeks of recovery, n=4) (**Extended Data Fig. 1e**). By comparing the transcriptome of old *versus* young samples, we identified a list of significantly differentially expressed genes (aging-DEGs, n=217) (**Supplementary Table 4**). PCA analysis of these aging-associated DEGs clearly separated old and young samples as expected. Remarkably, PCA placed old-OSKM profiles between old and young ones, suggesting that transient OSKM activation partially restores aging-related transcriptomic changes (**Fig. 2a, Supplementary Information**). To visualize the behavior of individual genes, we plotted the aging-DEGs and we colored those genes that were affected by OSKM. Interestingly, the large majority of genes upregulated by OSKM in old mice (pink dots) corresponded to genes with reduced expression upon aging (**Fig. 2b**). Conversely, those genes downregulated by OSKM in old mice (blue dots) were genes with upregulated expression upon aging (**Fig. 2b**). Looking then at the whole transcriptome, we first interrogated gene-sets well-established to change with aging, such as mTOR^24^ and DNA replication^25^. As expected, the mTOR signaling gene-set was upregulated in old *versus* young control mice (**Fig. 2c**), whereas the DNA replication gene-set was downregulated (**Fig. 2d**). Notably, old-OSKM samples behaved like young samples when compared to untreated old control mice (**Fig. 2c-d, Supplementary Table 5**). To better analyze the behavior of gene-sets across the three groups of samples, we performed a pattern analysis based on a Normal-Normal hierarchical model (gaga)^26^ to identify those gene-sets that (i) change significantly between young and old samples, and (ii) are similarly expressed in young and old-OSKM samples. This analysis identified a total of 179 gene-sets rejuvenated by OSKM (see pattern 1 in **Supplementary Table 6**). When the same analysis was performed after randomizing the samples (*i.e.* samples were randomly assigned to the three experimental groups: young, old and old-OSKM), only 44 gene-sets were found (see pattern 1 in **Supplementary Table 7**). More importantly, the gene-sets rejuvenated by OSKM included important aging-related processes, such as mTOR upregulation, insulin increase, reduction of NADPH and pyrimidine synthesis, and decline of mitochondrial processes, such as fatty acid oxidation, tricarboxylic acid cycle and oxidative phosphorylation^24, 27, 28^ (**Fig. 2e, Extended Data Fig. 2b-f**). All these gene-set alterations were ameliorated in old-OSKM mice (**Fig. 2e, Extended Data Fig. 2b-f**). DNA replication and repair are known to be reduced with aging^29, 30^. We observed that gene-sets related to the minichromosome maintenance (MCM) helicase, the DNA replication machinery, mismatch repair and base excision repair, were all reduced in our old mice and upregulated to young levels in old-OSKM mice (**Fig. 2f, Extended Data Fig. 2g**). Another important feature of aging is impaired protein homeostasis^31^. Again, this process was improved in old-OSKM mice (**Fig. 2g, Extended Data Fig. 2h**). Finally, loss of collagens occurs with age in pancreas^32^ and, remarkably, old-OSKM mice increased their levels of collagens (**Fig. 2h, Extended Data Fig. 2i**). Overall, transient OSKM activation appears to orchestrate a positive reconfiguration of the transcriptome against key hallmarks of aging.

**Figure 2.**
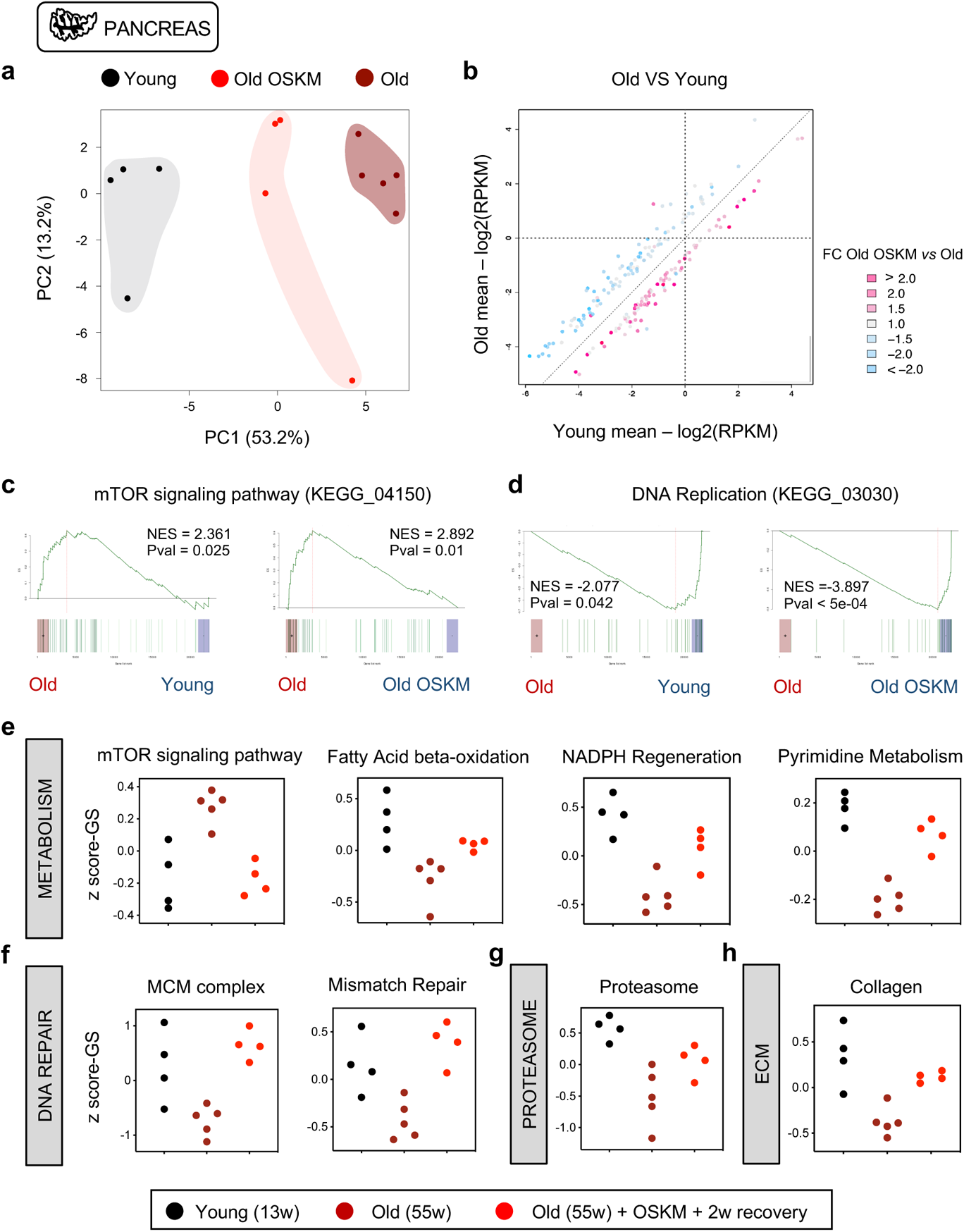
Transient OSKM reprogramming rejuvenates the transcriptome of old pancreas. **a,** Principal Component Analysis (PCA) of aging-related differentially expressed genes (DEGs: fold change>1.5 and raw pval<0.01) including young, old and old-OSKM pancreas. **b,** Representation of these aging-DEGs (217 genes in total) colored by their alteration of expression induced by OSKM: OSKM-upregulated genes are depicted in pink and OSKM-downregulated genes are depicted in blue. **c,** Enrichment analysis based on ROAST^52, 54^ was performed comparing young *versus* old, and old *versus* old-OSKM pancreas depicting: c, mTOR signaling pathway (KEGG_04150) and **d,** DNA Replication (KEGG_03030). Statistical significance in c and d was evaluated using Komolgorov-Smirnov test. **e-h,** Z-score representation of the expression profile with GS-adjustment for the indicated gene-sets among young, old and old-OSKM pancreas. Gene-sets have been selected for following this pattern: Young ≠ Old & Young= Old OSKM after a Normal-Normal hierarchical model (gaga)^26^ (5% FDR).

### Evidence of rejuvenation in tissues with low reprogramming susceptibility

Given its rejuvenating potential at the epigenetic and transcriptomic levels in the pancreas, we wondered whether other tissues that are less susceptible to reprogramming would nevertheless benefit from a single period of transient OSKM activation. For this, we performed a similar methylation profiling by RRBS analysis in the liver (5.3% of the promoters and 3.5% of the enhancers were covered) and spleen (25% of the promoters and 22.4% of the enhancers were covered), two tissues that modestly upregulate OSKM expression upon doxycycline treatment (**Extended Data Fig. 1c**). As before, we identified a list of DM promoters and enhancers during aging. Similar to the pancreas, PCA analysis of the methylation patterns of liver DM promoters and enhancers indicated that old-OSKM mice partially recovered a younger methylation pattern although the separation between experimental groups was not as clear as in the case of the pancreas (**Fig. 3a, Supplementary Information**). From the total combined number of aging-DM promoters and enhancers (n=108), a substantial fraction (61%) underwent rejuvenation (**Fig. 3b, Extended Data Fig. 3a-f, Supplementary Table 8**). This included *Foxa3* which serves as a pioneer transcription factor for the maintenance of liver-specific transcription^33^ (**Fig. 3b**). Other notable promoters were those of *Hoxd10*, a tumour suppressor gene whose promoter hypermethylation has been linked to hepatocellular carcinoma^34^, and *Thy1* whose expression stimulates liver regeneration^35^ (**Fig. 3b**). In the pancreas, we observed a temporal pattern for OSKM-induced methylation changes (remethylation during OSKM expression and demethylation during the recovery period). However, this temporal pattern was not evident in liver (**Extended Figure 3g**).

**Figure 3.**
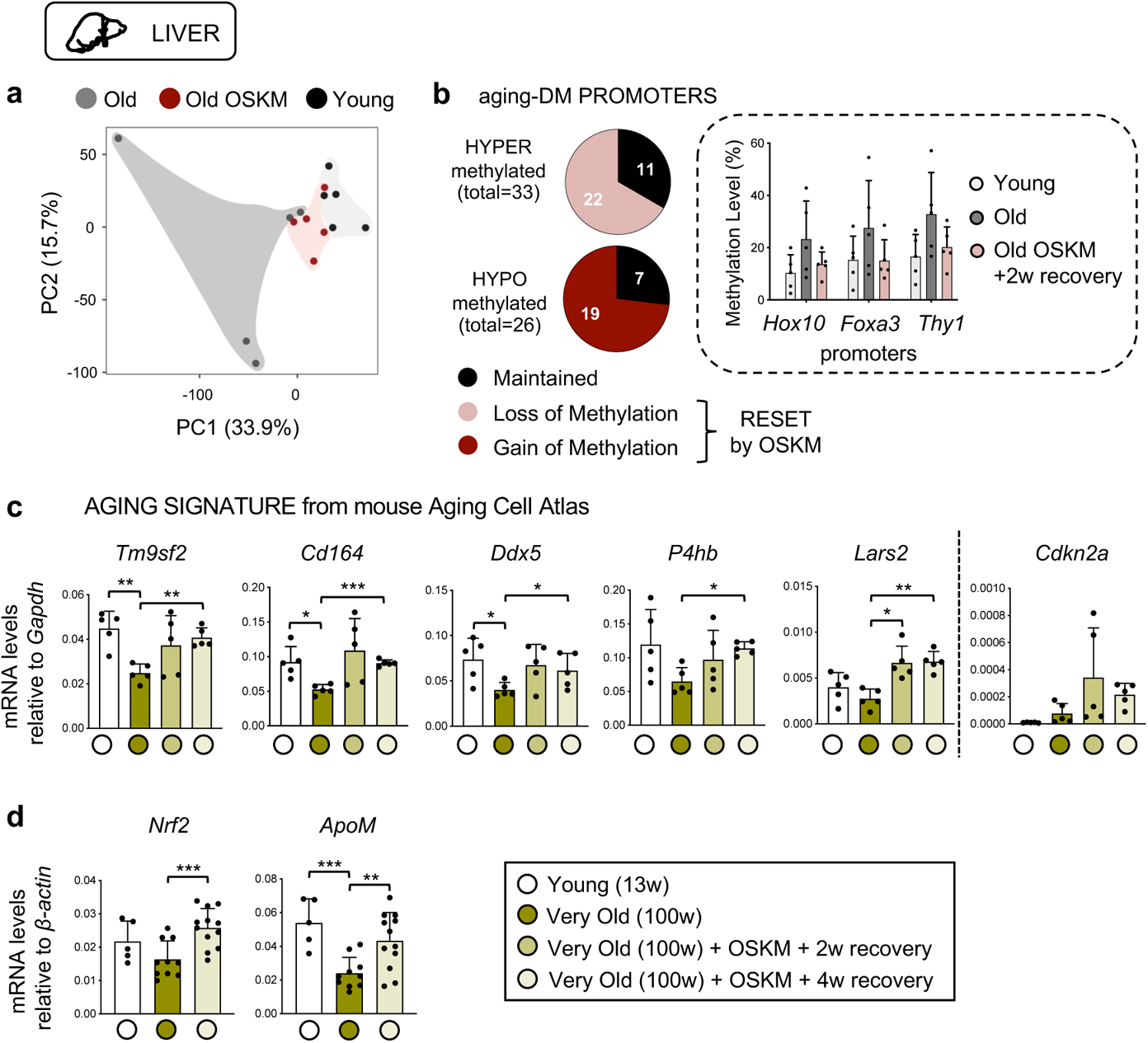
Old livers present rejuvenated features after transient OSKM reprogramming. **a,** Principal Component Analysis (PCA) of aging-related differentially methylated (DM) promoters including young, old and old-OSKM livers. **b,** DM promoters are divided into hyper- and hypomethylated during aging, and shown is the number of these promoters that alter their methylation profile due to transient OSKM activation. *Hox10, Foxa3* and *Thy1* promoters are representative examples of age-associated hypermethylated promoters that become demethylated in old OSKM livers. **c,** The expression of global aging genes identified by mouse Aging Cell Atlas^37^ was evaluated in very old livers (100 weeks; group 1 consists of 5 wild-type mice as control, 5 reprogrammable mice activating OSKM for 1 week and recovering for 2 weeks, and 5 reprogrammable mice activating OSKM for 1 week and recovering for 4 weeks) compared to young (13 weeks; n= 5) control livers. **d,** The expression of *Nrf2* and *ApoM*, two aging-associated genes, was measured in very old livers (100 weeks; group 1 as above together with group 2 consisting of other 5 wild-type mice as control and 7 reprogrammable mice activating OSKM for 1 week and recovering for 4 weeks). Statistical significance was evaluated using one-way ANOVA with Tukey’s multiple comparison method, and comparisons are indicated as *P < 0.05, **P < 0.01 and ***P < 0.001.

To obtain further insights into the aging-associated transcriptome of old-OSKM livers, we examined the top genes of a recently reported aging signature based on the Mouse Aging Cell Atlas^36^ that applies to multiple tissues including the liver and spleen but not to the pancreas^37^. In a pilot test, we measured by qRT-PCR a total of 15 mRNAs from this signature in young and old liver. Five of these genes were confirmed to be downregulated in the liver of our old mice (**Fig. 3c**). Interestingly, all these five genes recovered young levels of expression in the liver of old-OSKM mice (**Fig. 3c**). These findings were further validated in an independent cohort of very old mice (100 weeks) (**Extended Data Fig. 3h**). Finally, we tested the levels of two other genes associated to aging, namely, *Nrf2,* a key regulator of cellular redox homeostasis^38^, and *ApoM*, a high density lipoprotein (HDL) that promotes vascular homeostasis^39^. We confirmed that both genes are downregulated with aging in the liver of our very old mice, and this was reverted upon one cycle of OSKM (**Fig. 3d**). Notably, the levels of aspartate aminotransferase (AST/GOT) and alanine aminotransferase (ALT/GPT) in the serum of OSKM mice were significantly lower after 1 week of OSKM activation and 2 weeks of recovery reflecting a better liver function.

Senescent cells increase with aging and may account for up to 17% of the hepatocytes in extremely old mice (~3 years of age)^40^. Cellular senescence is a potent barrier for reprogramming^41^ and we wondered if our transient reprogramming protocol could reduce cellular senescence *in vivo*. For this, we examined the senescence marker p16^INK4a^ (*Cdkn2a*) and the senescence-associated cytokines *Mcp1* and *Cxcl2*, known to increase with aging in mouse liver^42^. As expected, all of these were increased with aging; however, their levels did not decline in any of the two cohorts of very old-OSKM mice (**Fig. 3c, Extended Data Fig. 3h,i**). The levels of p21^CIP1^ expression (*Cdkn1a*) were not informative because their levels did not change with aging nor with OSKM (**Extended Data Fig. 3h,i**). We also measured the number of γH2AX positive cells, a marker of *in vivo* senescence finding that the levels in the liver were similar in very old mice with or without 1 cycle of OSKM (**Extended Data Fig. 3k**). These observations suggest that a single period of transient OSKM *in vivo* may not be sufficient to rejuvenate the transcriptome of the senescent cells present in the aged liver.

In the spleen, PCA analysis of aging-DM promoters and enhancers did not allow to infer an epigenetic rejuvenation of this tissue (**Extended Data Fig. 4a,e, Supplementary Information**), although up to 163 promoter and enhancer regions showed evidence of rejuvenation according to their average methylation level (**Extended Data Fig. 4b-d, f-h, Supplementary Table 9**). To further explore the possibility of epigenetic rejuvenation in hematopoietic cells, we focused on the methylation of a specific intragenic region of the *Hsf4* gene^43–45^. First, we measured by pyrosequencing the methylation levels of the *Hsf4* region in the blood of mice of different ages, confirming a linear change of large magnitude, from ~20% methylation in young mice (~10 weeks) to ~55% methylation in very old mice (~100 weeks) (**Fig. 4a**). Then, we measured *Hsf4* methylation in the blood of very old mice, a group of reprogrammable mice and a control group of non-reprogrammable littermates, during a lifespan period of 5 weeks (1 week with doxycycline and 4 weeks without). During this time, it was possible to detect an increase in the *Hsf4* methylation levels of 4% in control mice (**Fig. 4b**). In contrast, reprogrammable mice reduced their average methylation levels by ~4% (**Fig. 4b**).

**Figure 4.**
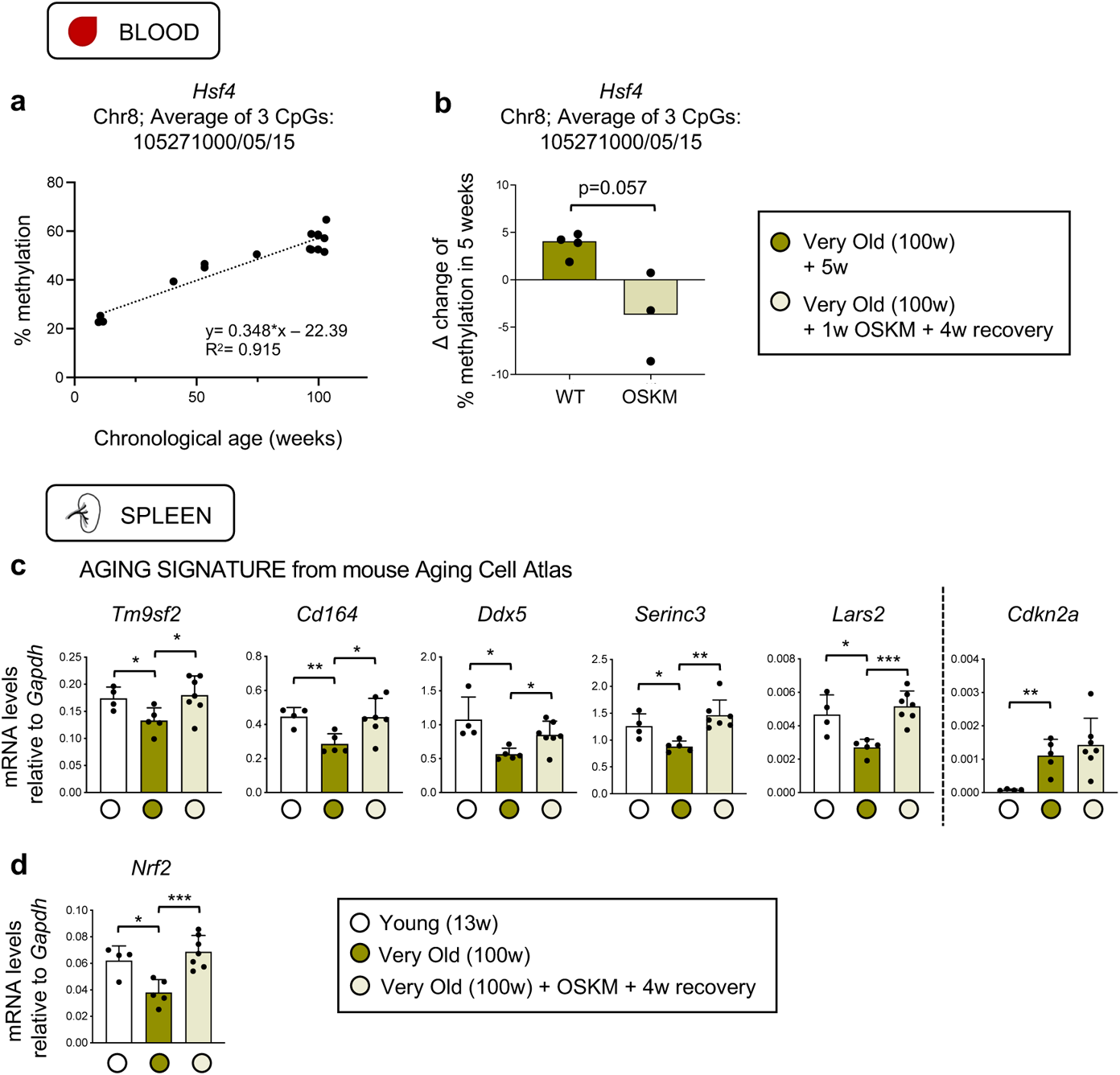
Evidences of OSKM-induced rejuvenation in haemopoietic cells. **a,** Correlation of the average methylation of 3 close CpGs located in the intragenic region of *Hsf4* gene (Chr8; at the positions 105271000,105271005 and 105271015), as measured in the blood of mice, with their chronological age (in weeks) of the mice. **b,** Δ change of the methylation levels in these CpGs of *Hsf4* in a period of 5 weeks comparing very old (100 weeks) wild-type mice (n=4) and reprogrammable mice (n=3) treated for 1 week with doxycycline and recovered for 4 weeks. Statistical significance was evaluated using Mann Whitney nonparametric t-test. **c,** The expression of global aging genes identified by mouse Aging Cell Atlas^37^ was evaluated in very old spleens (100 weeks; 5 wild-type mice as control, 7 reprogrammable mice activating OSKM for 1 week and 4 weeks of recovery) compared to young (13 weeks; n= 4) control spleens. **d,** The expression of *Nrf2*, an aging-associated gene, was measured in the same group of mice. Statistical significance was evaluated using one-way ANOVA with Tukey’s multiple comparison method, and comparisons are indicated as *P < 0.05, **P < 0.01 and ***P < 0.001.

We also tested the aging signature based on the Mouse Aging Cell Atlas in the spleen^37^. Interestingly, OSKM transient expression rescued the age-associated decline of seven of these genes while the senescent marker p16^INK4a^ (*Cdkn2a*) was unchanged (**Fig. 4c, Extended Data Fig. 4i**). Finally, *Nrf2* also declined in very old control spleens and its expression was reset to young levels in very old-OSKM spleens (**Fig. 4d**).

We conclude that a single cycle of OSKM expression has a detectable rejuvenating effect on tissues, such as liver, spleen and blood, that do not express high levels of OSKM and do not manifest obvious signs of histological alterations. Conceivably, some of the observed effects on liver, spleen and blood could be secondary to the direct rejuvenating actions of OSKM in other tissues, such as the pancreas.

### Serum metabolomic profiling reveals systemic benefits

The aforementioned results indicating epigenetic and transcriptional rejuvenation in several tissues prompted us to identify possible systemic signs of rejuvenation in the serum. To address this, mass spectrometric metabolomic analyses were performed on the sera of female mice (to reduce sex-related variations) from two independent experiments (each experiment analyzed separately by mass spectrometry). Each experiment included a group of young mice and a group of very old (~100 weeks) reprogrammable mice. The sera of very old reprogrammable mice were analyzed longitudinally, *i.e.* before and after a single cycle of reprogramming (1 week of doxycycline and 2 or 4 weeks of recovery). A total of 23 metabolites were identified as significantly changed between young and very old mice (**Fig. Extended Data Fig. 5a,b, Supplementary Table 10**). Out of these aging-associated metabolites, 4 were reversed after reprogramming (**Fig. 5a**). These metabolites are 4-hydroxyproline, thymine, trimethyl-lysine and indole-3-propionic acid, and some of them have been previously linked to aging. Serum 4-hydroxyproline has been previously reported to decline with aging^46^. This modified amino acid amounts to 14% of all the collagen and its presence in the serum is considered to reflect total collagen levels^47^. Notably, our transcriptomic analyses identified collagen synthesis as a gene-set downregulated with aging and rescued by OSKM in the pancreas (**Fig. 2h**). Thymine has been found to extend lifespan in *C. elegans*^48^. The other two metabolites, trimethyl-lysine and indole-3-propionic acid, have not been studied in the context of aging.

**Figure 5.**
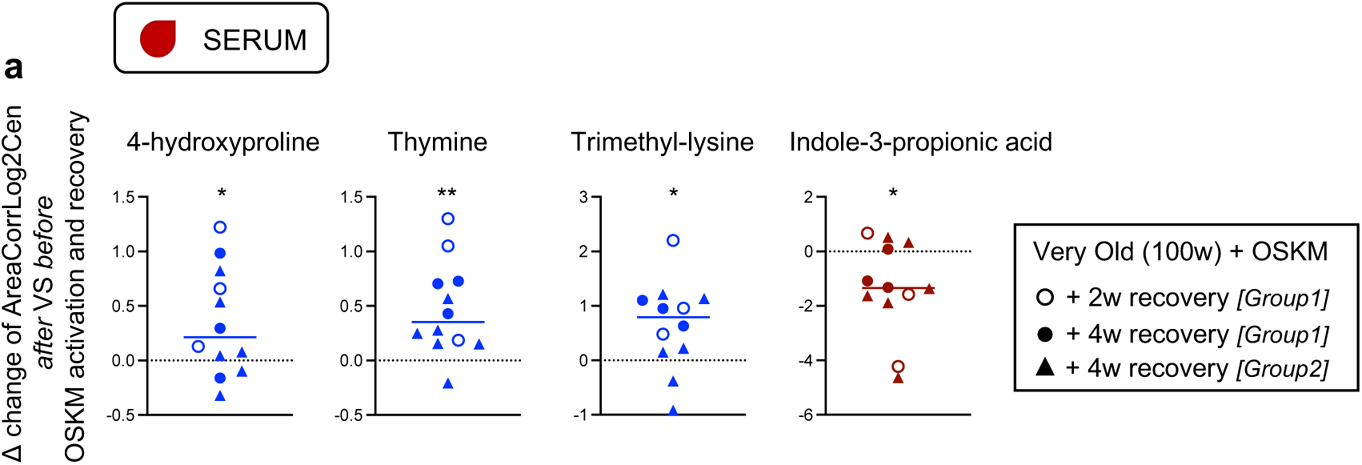
Metabolomic analysis in the serum of very old mice reveals systemic beneficial effects upon transient OSKM activation. **a,** Metabolomic analyses were performed on the sera of very old (100 weeks) female reprogrammable mice from two independent experiments (each experiment analyzed separately by mass spectrometry); *Group 1:* n=6 and *Group 2:* n=6. The sera of these mice were analyzed longitudinally, *i.e.* before and after a single cycle of reprogramming. The Δ change of the levels of 4 metabolites that were identified to change with aging, being either upregulated (4-hydroxyproline, thymine, trimethyl-lysine) or downregulated (indole-3-propionic acid), are depicted after activating OSKM for 1 week and recovering for 2 o 4 weeks. Statistical significance was evaluated using a paired t-test as values were confirmed to follow a normal distribution using the Shapiro-Wilk test. Comparisons are indicated as *P < 0.05, **P < 0.01 and ***P < 0.001.

## Discussion

In this work, we report that a single cycle of transient OSKM expression in naturally aged mice can elicit epigenetic, transcriptomic and serum metabolomic changes that partially restore younger patterns. We have focused on the effects of a single cycle of OSKM expression. The idea behind this choice was to identify and quantify early events induced by a single cycle of OSKM expression. Moreover, we have used a relatively low dose of doxycycline to avoid dramatic changes in cell identity thereby minimizing the risk of teratoma formation. This protocol of induction was sufficient to cause histologically detectable alterations in the pancreas, but not in the spleen or liver. Another relevant aspect of our experimental design is that we allowed for a recovery period of 2 to 4 weeks post-OSKM expression. This recovery period was sufficient for a full restoration of normal histology in the pancreas. Conceivably, the recovery period may capture those OSKM-induced changes that are more stable and, as we will argued below, it can be relevant for rejuvenation events secondary to OSKM expression.

In the case of the pancreas, we assessed whether methylation changes in aging-associated DMRs were already present at the time of switching off OSKM or rather acquired during the 2-weeks recovery period. First of all, about half of the changes observed at the time of switching off OSKM disappeared after 2 weeks. This illustrates the importance of allowing a recovery period to identify durable changes. Secondly, we observed interesting differences in the behavior of aging-DMRs depending on whether OSKM induced losses or gains of DNA methylation. The majority of gain-of-methylation events observed after recovery were already present at the time of extinguishing OSKM expression. In contrast, the majority of loss-of-methylation events were absent after OSKM expression and, therefore, were acquired during the recovery period. In line with these observations, pyrosequencing measurement of individual aging-methylated CpGs consistently showed more demethylation 4 weeks post-OSKM expression compared to 2 weeks. Together, we speculate that gains of methylation are closely linked to OSKM expression, whereas demethylation events occur secondary to OSKM expression during reversion. Conceivably, the reversion phase may be as important as the OSKM expression phase for molecular and cellular rejuvenation.

Analysis of the global transcriptome of the pancreas also allowed us to select for aging-related differentially expressed genes (aging-DEGs). Similar to the methylation data, one cycle of OSKM (1 week of induction and 2 weeks of recovery) was sufficient reset the levels of aging-DEGs to younger state. We have not been able to establish associations between specific aging-DMRs and the mRNA levels of the associated genes. This lack of association between aging-DMRs and transcription is, however, a general finding made by multiple researchers studying aging-DMRs (see detailed discussion in ^7^). For example, aging-DMRs often occur in bivalent, silent, promoters where gain of methylation does not impact the already silent transcriptional state^7^.

In the case of the spleen and liver, the methylation changes observed in aging-DMRs were not as clear as in the case of the pancreas, as judged by the principal component analyses. This is not entirely surprising if we consider the mild expression of the OSKM transgene in these tissues. Nevertheless, spleen and liver did show evidence of rejuvenation of a transcriptional signature of aging derived from the Mouse Aging Cell Atlas^37^. It seems, therefore, that transcriptional rejuvenation may be easier to achieve compared to epigenetic rejuvenation. Another layer of complexity comes from the fact that the rejuvenation features observed in a given tissue may reflect in part indirect effects caused by extra-tissular components, such as, serum factors or cells of hematological origin.

To evaluate molecular features of aging in the blood, we have focused on three closely located CpGs in an exon of the *Hsf4* gene that have called the attention of at least three independent groups^43–45^. Consistent with a previous report^43^, we found that the degree of methylation of these sites in peripheral blood is tightly linked to the age of the mice. We have measured the methylation levels of *Hsf4* in the blood of very old mice (~100 weeks of age) before and after a period of 5 weeks. In the case of control mice, we observed an increase in methylation during this period of 5 weeks. This was in contrast to mice exposed to one cycle of reprogramming (1 week of OSKM expression followed by 4 weeks of recovery) in which *Hsf4* methylation was decreased or maintained.

As a readout of systemic rejuvenation, we focused on serum metabolites. We analyzed the serum metabolome in two cohorts of very old mice (~100 weeks of age) in a longitudinal design, *i.e.* serum was analyzed before treatment and after one cycle of OSKM expression (1 week of doxycycline followed by 4 weeks of recovery). We have detected four serum metabolites that changed with aging and were reversed by OSKM in the two cohorts of old mice: 4-hydroxyproline, thymine, trimethyl-lysine and indole-3-propionic acid. Serum 4-hydroxyproline has been previously reported to decline with aging^46^ and it is considered to reflect total collagen content in the organism^47^. Of note, the transcriptional network for collagen synthesis was downregulated with aging and rescued by OSKM in the pancreas (**Fig. 2h**). Thymine has been found to extend lifespan in *C. elegans*^48^ and this is consistent with the aging-associated reduction that we have observed for this metabolite and its reversion by OSKM. Little is known about trimethyl-lysine and indole-3-propionic acid in connection with aging. Trimethyl-lysine is a modification abundant in chromatin and we speculate that its presence in the serum may derive from neutrophil-derived extracellular traps (NETs), a process by which neutrophils extrude their chromatin into the bloodstream^49^. Interestingly, the production of NETs by stimulated neutrophils declines with aging^49^, which would be consistent with a reduction in trimethyl-lysine.

In this work, we report stable reversion of molecular features of natural aging by a single cycle of OSKM expression. The observed rejuvenation features include DNA methylation, transcription and serum metabolites. These molecular manifestations of rejuvenation likely reflect a complex mixture of direct and indirect effects on multiple tissues. We hope this serves as the basis for future studies to dissect the mechanisms underlying OSKM-driven rejuvenation *in vivo*. Also, it may provide benchmarking to recapitulate OSKM-like rejuvenation with pharmacological or nutritional interventions.

## MATERIALS & METHODS

### Animal Procedures

Animal experimentation was performed between two different institutes: at the Spanish National Cancer Research Centre CNIO in Madrid and at the Institute of Research in Biomedicine IRB in Barcelona, according to protocols approved by the CNIO-ISCIII Ethical Committee for Research and Animal Welfare (CEIyBA) in Madrid, and by the Animal Care and Use Ethical Committee of animal experimentation of Barcelona Science Park (CEEAPCB) and the Catalan Government in Barcelona. We used the reprogrammable mice known as i4F-B which carries a ubiquitous doxycycline-inducible OSKM transgene, abbreviated as i4F, and inserted into the Pparg gene^11^. Mice of both sexes were used, and of different ages; young (females, 13 weeks), old (females, 55 weeks) and very old (males and females, 100 weeks). 0.2 mg/ml of Doxycycline hyclate BioChemica (PanReac) was administered in the drinking water supplemented with 7.5% sucrose for a period of 7 days. Mice were sacrificed two or four weeks after doxycycline removal. In the case of the intraperitoneal injection of doxycycline, young mice received 2.5 mg of doxycycline dissolved in 100μl of saline and sacrificed 24 hours later.

### DNA isolation

All pancreas, liver and spleen tissues were snap-frozen directly after collection. Genomic DNA was extracted from frozen tissues using the DNeasy Blood & Tissue Kit (Qiagen). In the case of blood samples, genomic DNA isolation was conducted according to a standard phenol-chloroform extraction protocol after red blood cell lysing. All samples were collected, processed for their methylation status and further analyzed.

### Reduced Representation Bisulfite Sequencing (RRBS) library preparation

RRBS libraries were prepared as described previously^18^. 500ng of genomic DNA was digested with MspI (Thermo Scientific). Following digestion, DNA fragments were end-repaired and T-tailed with Klenow fragment lacking 5’ → 3’ and 3’ → 5’ exonuclease activity (NEB). In-house adapters containing 8-nucleotide unique molecular identifier (UMI) sequences were ligated onto the DNA fragments with T4 DNA ligase (NEB). Excess adapters were removed by AMPure XP beads (Agencourt, 0.8X ratio). Libraries were bisulfite converted with the EZ-96 DNA Methylation-Direct MagPrep kit and cleaned up using an automated liquid handling platform (Agilent Bravo). The bisulfite converted libraries were amplified for 12 cycles with KAPA HiFi HotStart Uracil+ ReadyMix (Roche). The amplified libraries were cleaned up with AMPure beads (0.8X ratio) and quality checked using an Agilent High Sensitivity DNA kit on an Agilent 2100 Bioanalyzer system. Libraries passing quality checks were sequenced on an Illumina HiSeq 2500 sequencer for 75 bp paired-end sequencing. Reads were trimmed with Trim Galore (version 0.5.0). Trimmed reads were aligned to the mouse genome (GRCm38) with Bismark (version 0.20.0). Aligned reads were deduplicated based on UMI sequence and chromosomal position with Umibam (version 0.0.1). Bismark coverage files were generated from the deduplicated bam files using Bismark Methylation Extractor.

### RRBS DNA methylation analysis

Promoters were defined as −2000 bp to +500 bp of the transcription start site. Pancreas, spleen and liver enhancers were defined using published H3K27ac data^20, 21^. H3K27ac peaks were called using MACS and peaks that did not overlap promoter regions were classified as enhancers. Enhancers were linked to genes based on proximity. Enhancers were linked to the nearest gene within 1 Mb. The methylation analysis was carried out in Seqmonk. CpG sites were only carried forward in the analysis if they were covered by at last 5 reads in all the samples of that tissue type. The methylation level of each promoter and enhancer was calculated using the mean of all the methylation levels of the CpGs within those regions. Differential methylation analysis was carried out using logistic regression. Regions were identified as significant if their P-value was below 0.05 and if they demonstrated a minimum methylation difference of at least 10% DNA methylation. Rejuvenated regions were those in which the mean methylation in the old-OSKM group was closer to the mean of the young group than to the mean of the old group.

### Bisulfite pyrosequencing

DNA methylation patterns were validated by bisulfite pyrosequencing. Bisulfite modification of DNA was performed with the EZ DNA methylation-gold kit (Zymo Research) following manufacturer’s instructions. The sets of primers for PCR amplification and sequencing were designed using the specific software PyroMark assay design (version 2.0.01.15). Mm10 genome was used for the alignments. The primers used were: *1_133696298-311* forward primer: 5’-AGAGTGTTAGAGTTGGAGAGAT-3’, *1_133696298-311* reverse primer: 5’-[Btn]AAAAAAACCTCTAACCTCCATATATC-3’ (Sequencing: 5’-TGGAGAGATTTTGAAGTT-3’), *2_90532378* forward primer: 5’-GGTTTGGAATTTGGTTTATGTATAGA-3’, *2_90532378* reverse primer: 5’-[Btn]CTCAAACCAAAAAACCCTAATCTCC-3’ (Sequencing: 5’-TTTTTTTTAAGAGAATAGGATTATA-3’), *11_75813125* forward primer: 5’-GGGTTAGGTTGAGTTTTTTAGAATGAAT-3’, *11_75813125* reverse primer: 5’-[Btn]ACCCTCTCCATCTATACCTACTCC-3’ (Sequencing: 5’-AG-GGTTTTGTATTTAATTTTTATT-3’), *12_112670818* forward primer: 5’-AGAGGTTGGGATTGGTAAGGATT-3’, *12_112670818* reverse primer: 5’-[Btn]TTACCCCAAAACAAACATCTCACCC-3’ (Sequencing: 5’-GGTAAGGAT-TATTTTAGGGTT-3’), *14_121078071* forward primer: 5’-ATGAGATGATTTTAG-TTAAGGATTTAGTT-3’, *14_121078071* reverse primer: 5’-[Btn]AT-TACACAAAACTTCCAACTTACT-3’ (Sequencing: 5’-ATAGGAA-TAAAAGTTTTTTGATAA-3’), *14_75862317-319-323* forward primer: 5’-GAAGGAAGGGAATTTTGAGATTTG-3’, *14_75862317-319-323* reverse primer: 5’-[Btn]TCTCCCAAAACTATACCATCACCA-3’ (Sequencing: 5’-TTTGTTTTT-GTTTTTTTATTATAAG-3’), *19_34218524* forward primer: 5’-TGATTT-GGTTAAAGGTAGAAAAGTAAGA-3’, *19_34218524* reverse primer: 5’-[Btn]AACCTTTATAAACAATCAATAAATATACCT-3’ (Sequencing: 5’-GGTTTTT-GTGGATAAATATTA-3’), *19_43890303-291* forward primer: 5’-GGAT-TTAGGTGGGTTTTATTTAGAAAATG-3’, *19_43890303-291* reverse primer: 5’-[Btn]TCCCAATACCCACAATCCCTTTTT-3’ (Sequencing: 5’-GTTTTATTTA-GAAAATGGTTTGGA-3’), *19_58600208* forward primer: 5’-AGAGGAAA-TAATTTTATAGTGTAGGTAAGA-3’, *19_58600208* reverse primer: 5’-[Btn]CAAATTCTCAACCATAAAAATCACTCTA-3’ (Sequencing: 5’-TGGGTG-GAAATGTGA-3’), *mHsf4* forward primer: 5’-GTGYGTYGTAAGGTGGGATAAATTGTAGAAAAAATG-3’, *mHsf4* reverse primer: 5’-[Btn]TCCRTACTCTCCTACACTCCTCTCAAAACTTA-3’ (Sequencing: 5’-ATGGTGTTTTTTGTTTGTAG-3’). After PCR amplification, pyrosequencing and quantification were performed using PyroMark Q24 reagents, equipment, and software (Qiagen).

### RNA isolation and analysis of mRNA levels

Total RNA was extracted from pancreas samples using guanidine thiocyanate, followed by acid phenol-chloroform extraction. For the rest of the tissue samples (liver, spleen), total RNA was isolated with Trizol (Invitrogen), following provider’s recommendations. Up to 5 μg of total RNA was reverse transcribed into cDNA using iScriptTM Advanced cDNA Synthesis Kit for RT-qPCR (Bio-Rad). Quantitative real time-PCR was performed using GoTaq® qPCR Master Mix (Promega) in a QuantStudio 6 Flex thermocycler (Applied Biosystem) or 7900HT Fast (Applied Biosystem). For input normalization, we used the housekeeping genes *Gapdh*, *β-actin* and *18S.* The primers used were: *Gapdh* forward primer: 5’-TTCACCACCATGGAGAAGGC-3’, *Gapdh* reverse primer: 5’-CCCTTTT-GGCTCCACCCT-3’; *18S* forward primer: 5’-GTAACCCGTTGAACCCCATT-3’, *18S* reverse primer: 5’-CCATCCAATCGGTAGTAGCG-3’; *β-actin* forward primer: 5’-TGTTACCAACTGGGACGACA-3’, *β-actin* reverse primer: 5’-GGGGTGTTGAAGGTCTCAAA-3’; *Nanog* forward primer: 5’-CAAGGGTCTGCTACTGAGATGCTCTG-3’, *Nanog* reverse primer: 5’-TTTTGTTTGGGACTGGTAGAAGAATCAG-3’; *Oct4* (endogenous) forward primer: 5’-TCTTTCCACCAGGCCCCCGGCTC-3’, *Oct4* (endogenous) reverse primer: 5’-TGCGGGCGGACATGGGGAGATCC-3’; *Tfe3* forward primer: 5’-TGCGTCAGCAGCTTATGAGG-3’, *Tfe3* reverse primer: 5’-AGACACGCCAATCACAGAGAT-3’; *E2A-c-Myc* forward primer, 5-GGCTGGAGATGTTGAGAGCAA-3, *E2A-c-Myc* reverse primer 5-AAAGGAAATCCAGTGGCGC-3; *Tm9sf2* forward primer: 5’-GCAACGAGTGCAAGGCTGATA-3’, *Tm9sf2* reverse primer: 5’-CCCCGAATAATACCTGACCAAGA-3’; *Cd164* forward primer: 5’-GTGTTTCCTGTGTTAATGCCAC-3’, *Cd164* reverse primer: 5’-CACAAGTCAG-TGCGGTTCAC-3’; *Ddx5* forward primer: 5’-CGGGATCGAGGGTTTGGTG-3’, *Ddx5* reverse primer: 5’-GCAGCTCATCAAGATTCCACTTC-3’; *P4hb* forward primer: 5’-GCCGCAAAACTGAAGGCAG-3’, *P4hb* reverse primer: 5’-GGTAGCCACGGACACCATAC-3’; *Lars2* forward primer: 5’-CATAGAGAG-GAATTTGCACCCTG-3’, *Lars2* reverse primer: 5’-GCCAGTCCTGCTTCATA-GAGTTT-3’; *Serinc3* forward primer: 5′-GTCCCGTGCCTCTGTAGTG-3′, *Serinc3* reverse primer: 5′-CAAGACACAATAGTGCCAAGGAA-3′; *Nptn* forward primer: 5′-CGCTGCTCAGAACGAACCAA-3′, *Nptn* reverse primer: 5′-GCTGGAAGTGAGGTTACACTG-3′; *Cdkn2a* forward primer: 5’-CGAACTCTTTCGGTCGTACCC-3’, *Cdkn2a* reverse primer: 5’-CGAATCTGAACCGTAGTTGAGC-3’; *Cdkn1a* forward primer: 5’-TCTGAGCGGCCTGAAGATTC-3’, *Cdkn1a* reverse primer: 5’-CTGCGCTTGGAGTGATAGAA-3’; *Cxcl2* forward primer: 5′-CTCAAGGGCGGTCAAAAAGT-3′, *Cxcl2* reverse primer: 5′-TTTTTCTTTCTCTTTGGTTCTTCC-3′; *Mcp1* forward primer: 5′-ATTGG-GATCATCTTGCTGGT-3′, *Mcp1* reverse primer: 5′-CCTGCTGTTCACAGTT-GCC-3′; *Nrf2* forward primer: 5′-CTGAACTCCTGGACGGGACTA-3′, *Nrf2* reverse primer: 5′-CGGTGGGTCTCCGTAAATG-3′; *ApoM* forward primer: 5′-TAACTCCATGAATCAGTGCCCT-3′, *ApoM* reverse primer: 5′-CCCGCAATAAAGTACCACAGG-3′.

### Bulk RNA sequencing and analysis

#### RNA and libraries preparation

Total RNA was extracted from pancreas tissues as described above, and treated with the NEBNext rRNA depletion kit (New England Biolabs) to deplete the ribosomal RNA (rRNAs). For the library preparation, we used the NEBNext Ultra II RNA library prep kit for Illumina (New England Biolabs) (9 cycles of amplification). Sequencing was performed on a HiSeq 2500 Sequencing System of Illumina; the type of sequencing was 50 nt Paired-End and 55 million reads were obtained per sample.

#### Pre-processing

Paired-end reads were aligned to the mm10 genome UCSC Genome Browser using STAR 2.3.0e^50^ with the following parameter values: outFilterMismatchNoverLmax=0.05; outFilterMatchNmin=25; the rest of parameters were set to their default values. RPKM estimations per isoforms were computed with the R package Casper^51^ which were then aggregated at the gene level (entrez). Reads mapping to five or more locations were excluded (parameter keep.multihits set to FALSE). The resulting RPKM matrix was log2-transformed, quantile normalized and a-priori corrected by the total number of reads in each sample; for doing so, a linear model was fitted to the expression matrix gene-wise, in which the group condition was included as explanatory variable.

#### Functional enrichment analysis

Pathway enrichment analysis for group comparisons of gene expression was performed using a modification of ROAST^52^, a rotation-based approach implemented in the R package limma^53^ that is especially suitable for small size experiments and is based on limma differential expression. Such modifications were implemented to accommodate the re-standardized maxmean statistic in the ROAST algorithm^54^, in order to enable it for competitive testing^55^. For doing so, genes were annotated according to Gene Ontology (GO)^56^, Broad Hallmarks^57^ and Kegg^58^ gene sets collections. GO and Kegg terms were retrieved from R package org.Mm.eg.db^59^, while Broad Hallmark sets were translated to mouse homologous genes using the R package biomaRt^60^. For visualization and interpretation purposes, results were represented with Komolgorov-Smirnov based statistic usually used in Gene Set Enrichment Analysis (GSEA)^61^.

#### Pattern analysis at gene-set level

To better analyze the behavior of gene-sets across the three groups of samples, we used a hierarchical model based on a mixture of Normal distributions (Normal-Normal)^62^ as implemented in the gaga^26^ R package. This model allows to classify genes according to their expression pattern across the groups of interest. To get a measure of the pathway activity in the transcriptomic data, we summarized each pathway as a gene expression signature. For doing so, expression values were centered and scaled gene-wise according to the mean and the standard deviation computed across samples, which were then averaged across all genes included in a given gene-set. In addition, a global signature was computed using all the genes in the expression matrix and used for a-priori correction of gene-set scores by fitting a linear model, in which the group condition was included as explanatory variable. This strategy has proven to be useful to alleviate systematic biases due to the gene-correlation structure present in the data, and to adjust by the expectation under gene randomization, i.e., the association expected for a signature whose genes have been chosen at random^54, 63^. Pathway scores were merged with gene level expression and a gaga^26^ pattern analysis was performed using a Normal-Normal model fit, accordingly to their observed distribution. In these analyses, the threshold for statistical significance was set at 5% FDR. All statistical analyses were carried out using R and Bioconductor [M19].

### Metabolomics

#### Sample preparation serum

A volume of 25 µL of serum were mixed with 250 µL of a cold solvent mixture with ISTD (MeOH/Water/Chloroform, 9/1/1, −20°C), in a 1.5 mL microtube, before being vortexed and centrifugated (10 min at 15000 g, 4°C). The upper phase of the supernatant was split into three parts: 50 µL was used for GC-MS experiment in injection vial, 30 µL was used for the SCFA (Short Chain Fatty Acids) UHPLC-MS method, and 50 µL was used for other UHPLC-MS experiments^64^.

### Widely-targeted analysis of intracellular metabolites gas chromatography (GC) coupled to a triple quadrupole (QQQ) mass spectrometer

GC-MS/MS method was performed on a 7890B gas chromatograph (Agilent Technologies, Waldbronn, Germany) coupled to a triple quadrupole 7000C (Agilent Technologies, Waldbronn, Germany) equipped with a high sensitivity electronic impact source (EI) operating in positive mode^64^.

#### Targeted analysis of bile acids by ion pairing ultra-high performance liquid chromatography (UHPLC) coupled to a Triple Quadrupole (QQQ) mass spectrometer

Targeted analysis was performed on a RRLC 1260 system (Agilent Technologies, Waldbronn, Germany) coupled to a Triple Quadrupole 6410 (Agilent Technologies) equipped with an electrospray source operating in positive mode. Gas temperature was set to 325°C with a gas flow of 12 L/min. Capillary voltage was set to 4.5 kV^64^.

#### Targeted analysis of polyamines by ion pairing ultra-high performance liquid chromatography (UHPLC) coupled to a Triple Quadrupole (QQQ) mass spectrometer

Targeted analysis was performed on a RRLC 1260 system coupled to a Triple Quadrupole 6410 (Agilent Technologies) equipped with an electrospray source operating in positive mode. The gas temperature was set to 350°C with a gas flow of 12 l/min. The capillary voltage was set to 3.5 kV^64^.

#### Targeted analysis of Short Chain Fatty Acid by ion pairing ultra-high-performance liquid chromatography (UHPLC) coupled to a 6500+ QTRAP mass spectrometer

Targeted analysis was performed on a RRLC 1260 system coupled to a 6500+ QTRAP (Sciex, Darmstadt, Germany) equipped with an electrospray ion source^64^.

#### Untargeted analysis of intracellular metabolites by ultra-high performance liquid chromatography (UHPLC) coupled to a Q-Exactive mass spectrometer. Reversed phase acetonitrile method

The profiling experiment was performed with a Dionex Ultimate 3000 UHPLC system (Thermo Scientific) coupled to a Q-Exactive (Thermo Scientific) equipped with an electrospray source operating in both positive and negative mode and full scan mode from 100 to 1200 m/z. The Q-Exactive parameters were: sheath gas flow rate 55 au, auxiliary gas flow rate 15 au, spray voltage 3.3 kV, capillary temperature 300°C, S-Lens RF level 55 V. The mass spectrometer was calibrated with sodium acetate solution dedicated to low mass calibration^64^.

#### Differential expression in metabolomics data

To assess differences of metabolites levels in serum samples, a linear mixed-effect was fitted for each metabolite separately in which the biological specimen was included as a random effect. Wald tests derived from the models were used to assess statistical significance. These analyses were performed with R packages lme4^65^, lmerTest^66^ and multcomp^67^.

### Histological analysis

Tissue samples were fixed overnight in 10% neutral buffered formalin (4% formaldehyde in solution; Sigma), paraffin-embedded, sectioned at a thickness of 3 μm, mounted in Superfrost®plus slides and dried. Slides were deparaffinized in xylene and re-hydrated through a series of graded ethanol until water. Sections were stained with hematoxylin and eosin (HE) for the visualization of the tissue architecture. For immunohistochemistry, sections were stained with a Rabbit mAb Phospho-Histone H2A.X (Ser139) (20E3) (Cell Signaling, ref: 9718). Slides were dehydrated, cleared and mounted with Toluene-Free mounting medium for microscopic evaluation. Whole digital slides were acquired with a slide scanner and images captured with the NanoZoomer Digital Pathology software (NDP.view2).

### Statistical analysis

Mice were randomly allocated to their experimental groups, except from the cohort of very old mice (100 weeks) which were distributed according to their pre-determined type (mouse genotype) and therefore there was no randomization. Quantitative PCR data were obtained from independent biological replicates (n values correspond to the number of mice; technical replicates of PCR were not considered in the n value). Statistical analyses were carried out using GraphPad Prism v8.0 (GraphPad software) or as it is indicated in each specific method, and stated at the figure legends.

### Data availability

All sequencing data are deposited in GEO under the super-series accession number: GSE156558. To access the data, you can use the password: wnsvcagwbjwhbsp. All the rest of the data is available in the main text or the supplementary materials.

## ACKNOWLEDGMENTS

We thank assistance from IRB core facilities. D.C. was recipient of a fellowship from “laCaixa” Foundation. Work in the laboratory of M.F.F. was funded by FICYT (IDI/2018/146), Fundación de la AECC (PROYE18061FERN) and Fondo de Investigación Sanitaria (FIS) (PI18/01527). Work in the laboratory of G.K. was supported by the Ligue contre le Cancer (équipe labellisée); Agence National de la Recherche (ANR) – Projets blancs; ANR under the frame of E-Rare-2, the ERA-Net for Research on Rare Diseases; AMMICa US23/CNRS UMS3655; Association pour la recherche sur le cancer (ARC); Association “Ruban Rose”; Cancéropôle Ile-de-France; Chancelerie des universités de Paris (Legs Poix), Fondation pour la Recherche Médicale (FRM); a donation by Elior; European Research Area Network on Cardiovascular Diseases (ERA-CVD, MINOTAUR); Gustave Roussy Odyssea, the European Union Horizon 2020 Project Oncobiome; Fondation Carrefour; High-end Foreign Expert Program in China (GDW20171100085), Institut National du Cancer (INCa); Inserm (HTE); Institut Universitaire de France; LeDucq Foundation; the LabEx Immuno-Oncology (ANR-18-IDEX-0001); the RHU Torino Lumière; the Seerave Foundation; the SIRIC Stratified Oncology Cell DNA Repair and Tumor Immune Elimination (SOCRATE); and the SIRIC Cancer Research and Personalized Medicine (CARPEM). This study contributes to the IdEx Université de Paris ANR-18-IDEX-0001. Work in the laboratory of M.S. was funded by the IRB, and by grants from Spanish Ministry of Economy co-funded by European Regional Development Fund (ERDF) (SAF2013-48256-R), European Research Council (ERC-2014-AdG/669622), Secretaria d’Universitats i Recerca del Departament d’Empresa i Coneixement of Catalonia (Grup de Recerca consolidat 2017 SGR 282) and “laCaixa” Foundation.

## AUTHOR CONTRIBUTIONS

D.C. designed most experiments, performed animal experimentation, collected samples, performed RNA analysis, contributed to bioinformatic data analysis, prepared the figures, and co-wrote the manuscript. D.G. and W.R. designed, performed, analysed, and interpreted the RRBS methylome data. L.M. designed and performed some reprogramming experiments, collected tissue samples and extracted genomic DNA. R.G.U. and M.F.F. designed, performed, analysed, and interpreted the bisulfite pyrosequencing experiments. A.B. and C.S.O-A. designed and contributed to bioinformatic analysis. M.Abad performed the first rejuvenation studies and provided advise throughout the project. D.E.M.H contributed to the original conception and design of the study. M.Aguilera performed all the histological stainings. N.P. supervised and interpreted the pathological analyses, and blindly scored the histological stainings. S.D., F.A., N.N. and G.K. performed, analysed, and interpreted the metabolomics data. M.S. designed and supervised the study, analysed the data, and co-wrote the manuscript. All authors discussed the results and commented on the manuscript.

---

**Extended Data Figure 1.**
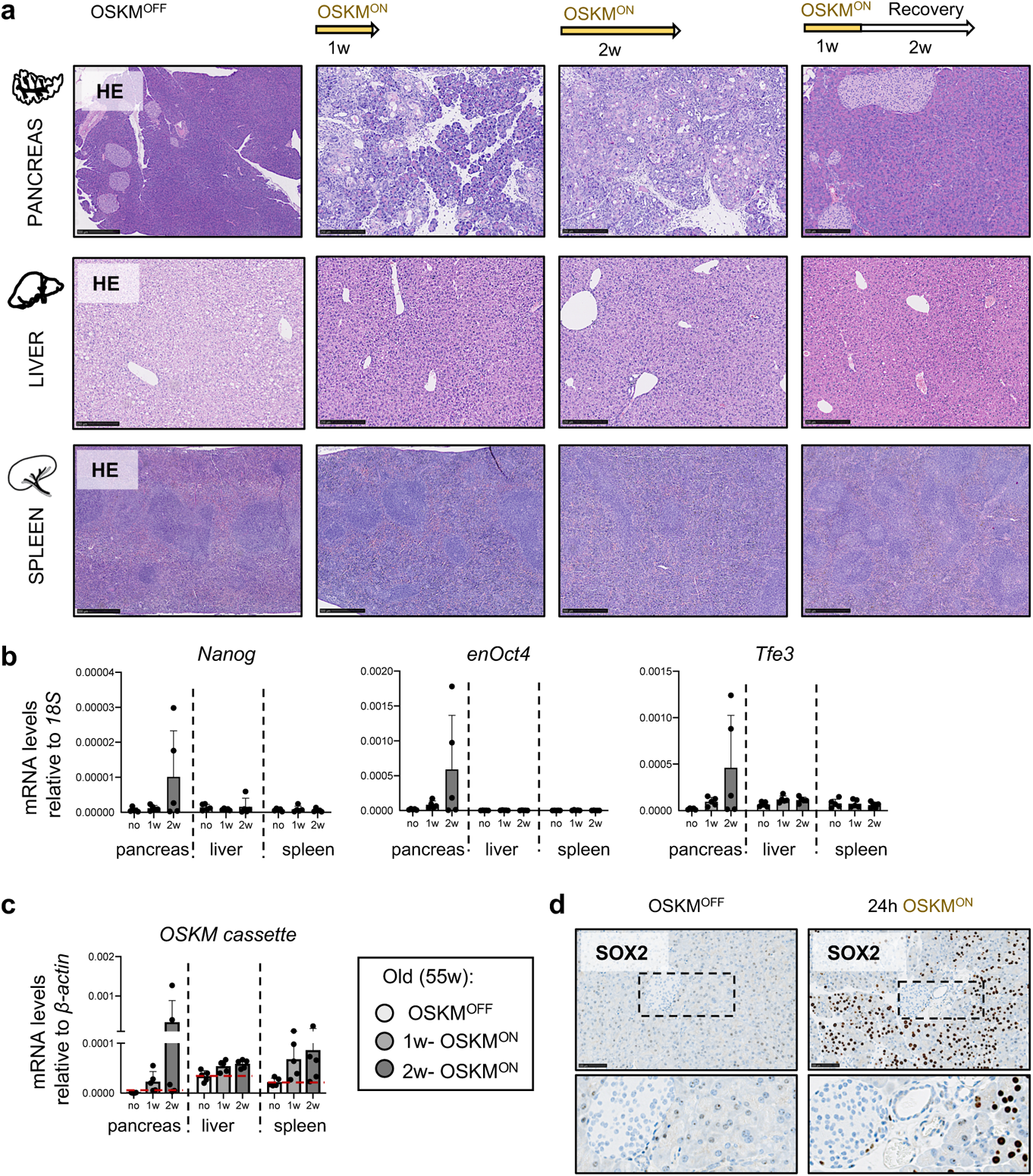
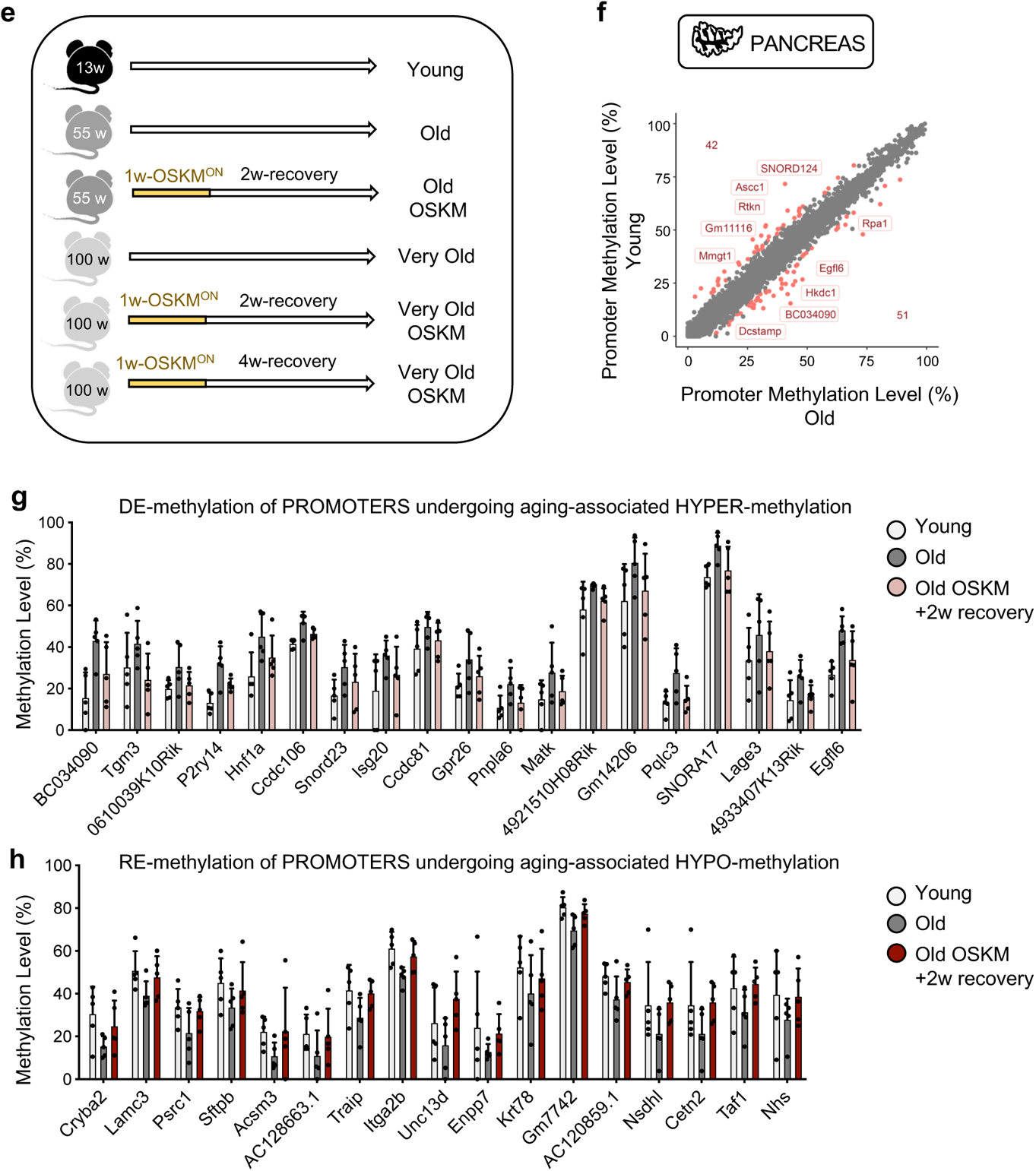
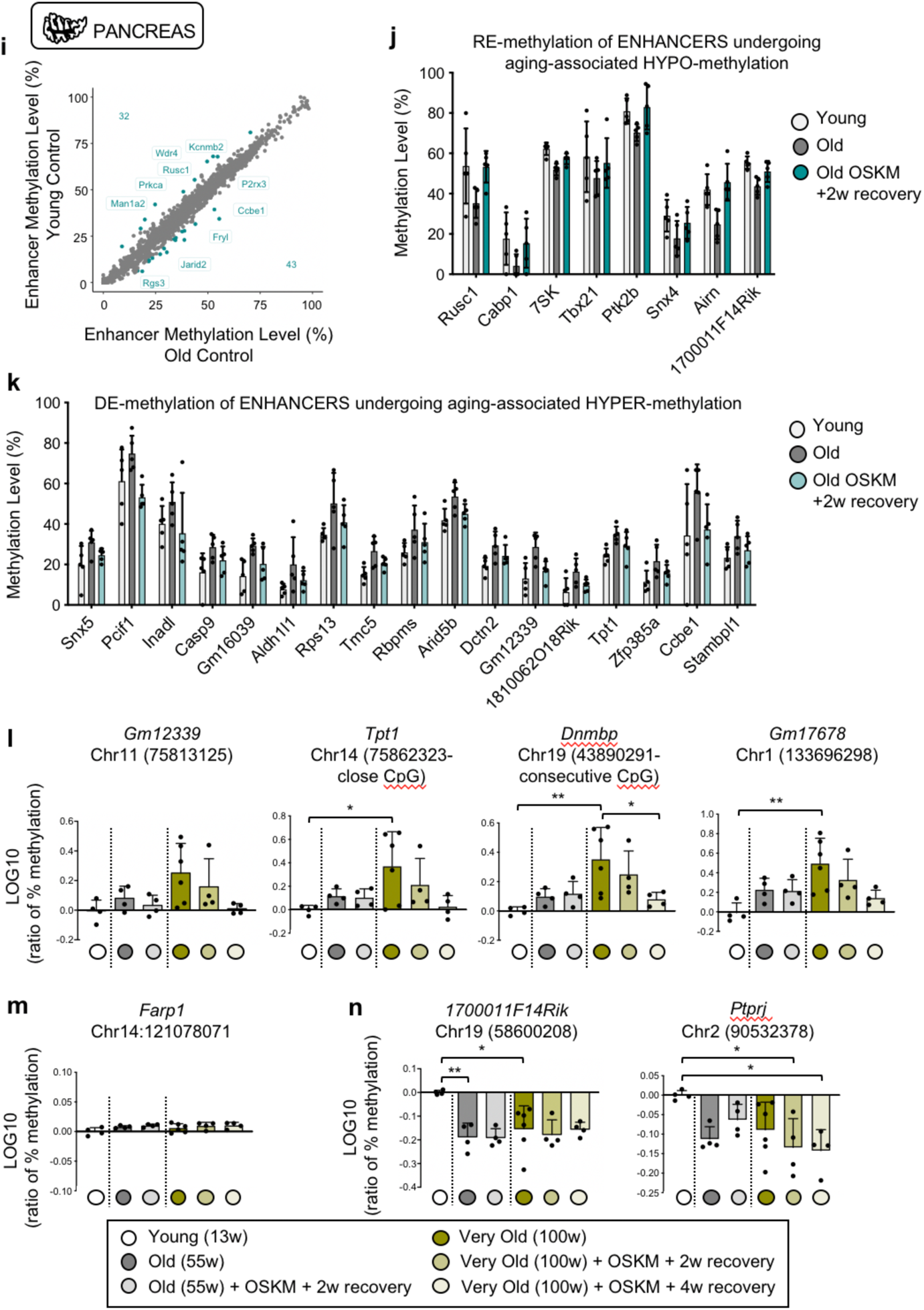
Methylation profile of aging-associated differentially methylated promoters and enhancers in old-OSKM pancreas. **a,** Hematoxylin Eosin (HE) of pancreas, liver and spleen of old (55 weeks) mice treated without or with doxycycline for one week, two weeks or one week followed by two weeks of recovery. **b,** RNA expression levels of pluripotency markers *Nanog*, endogenous *Oct4* (*enOct4*) and *Tfe3* in the indicated tissues (n=5 females). **c,** RNA expression levels of *OSKM cassette* using *E2A-c-Myc* primers in the same tissues. **d,** Immunohistochemistry of SOX2 in the pancreas of young reprogrammable mouse (10 weeks) 24 hours after intraperitoneal injection of doxycycline compared to untreated reprogrammable mouse. **e,** Schematic representation of the experimental groups: young (n=5 females, reprogrammable untreated), old (n=5 females, reprogrammable untreated), old-OSKM (n=5 females, reprogrammable treated with doxycycline), very old (n=6 males and females, wild-type treated with doxycycline), very old (n=4 males and females, reprogrammable treated with doxycycline plus 2 weeks recovery), very old-OSKM (n=4 males and females, reprogrammable treated with doxycycline plus 4 weeks recovery recovery). **f,** Identification of differentially methylated (DM) promoters in young *versus* old control pancreas. **g,** A set of gene promoters with decreased methylation levels undergoing aging-associated hypermethylation. **h,** A group of gene promoters with gain of methylation undergoing aging-associated hypomethylation. **i,** Identification of differentially methylated (DM) enhancers in young *versus* old control pancreas. **j,** A group of gene enhancers with gain of methylation undergoing aging-associated hypomethylation. **k,** A set of gene enhancers with decreased methylation levels undergoing aging-associated hypermethylation. **l,** Methylation levels of four CpGs, located in regions hypermethylated with aging measured and validated by bisulfite pyrosequencing and **m,** one non-validated CpG. **n,** Methylation levels measured by bisulfite pyrosequencing of two CpGs located in regions hypomethylated with aging.

**Extended Data Figure 2.**
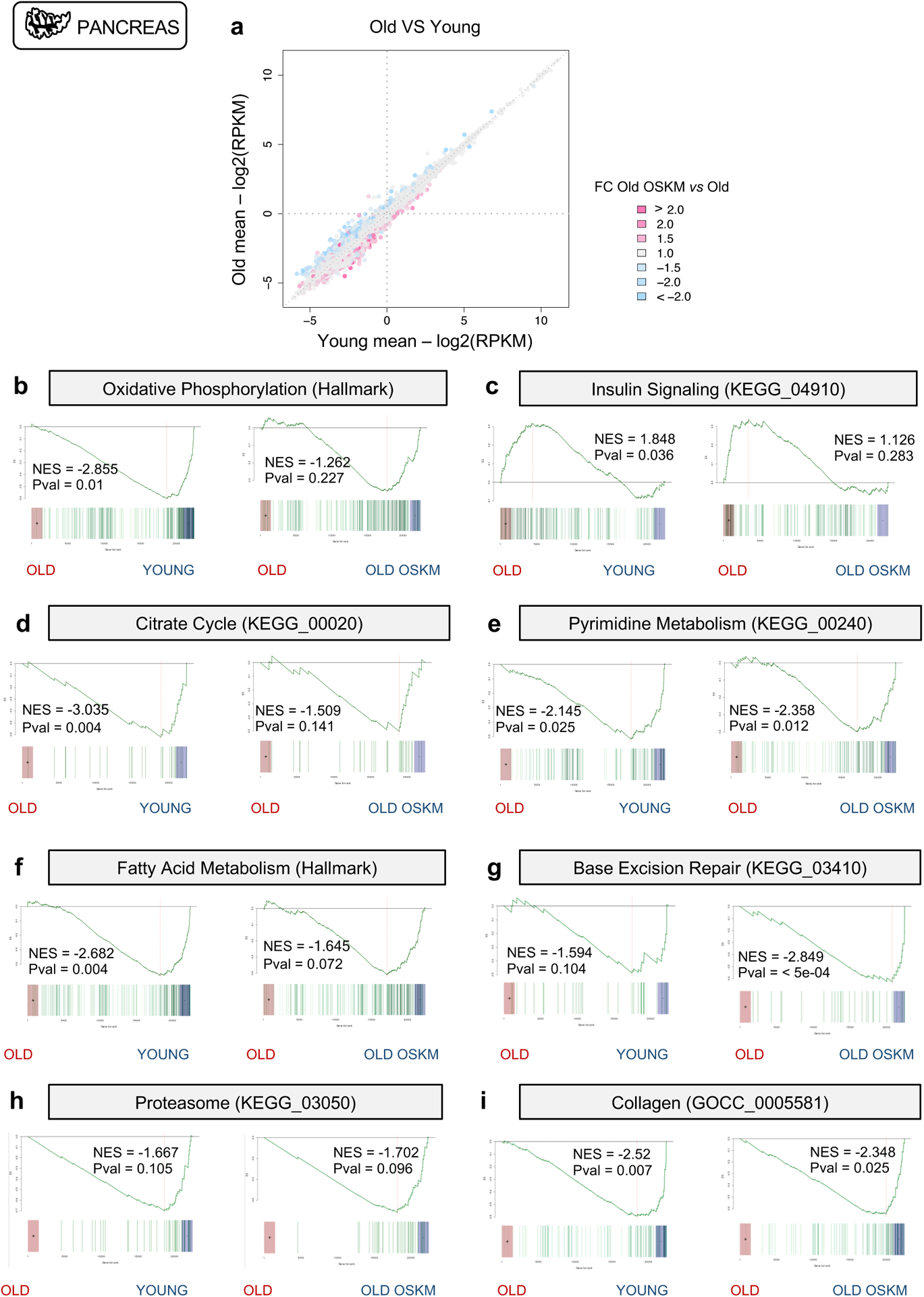
Transcriptional rejuvenation in old-OSKM pancreas. **a,** Representation of all DEGs in old *versus* young samples colored by their alteration of expression induced by OSKM: OSKM-upregulated genes are depicted in pink and OSKM-downregulated genes are depicted in blue. **b,** Enrichment analysis based on ROAST^52, 54^ was performed comparing young *versus* old, and old *versus* old-OSKM pancreas. Old mice are 55 weeks of age. The following processes are depicted: **b,** Oxidative Phosphorylation (Hallmark), **c,** Insulin Signaling (KEGG_04910), **d,** Citrate Cycle (KEGG_00020), **e,** Pyrimidine Metabolism (KEGG_00620), **f,** Fatty Acid Metabolism (Hallmark), **g,** Base Excision Repair (KEGG_03410), **h,** Proteasome (KEGG_03050), **i,** Collagen (GOCC_0005581). Statistical significance was evaluated using Komolgorov-Smirnov test.

**Extended Data Figure 3.**
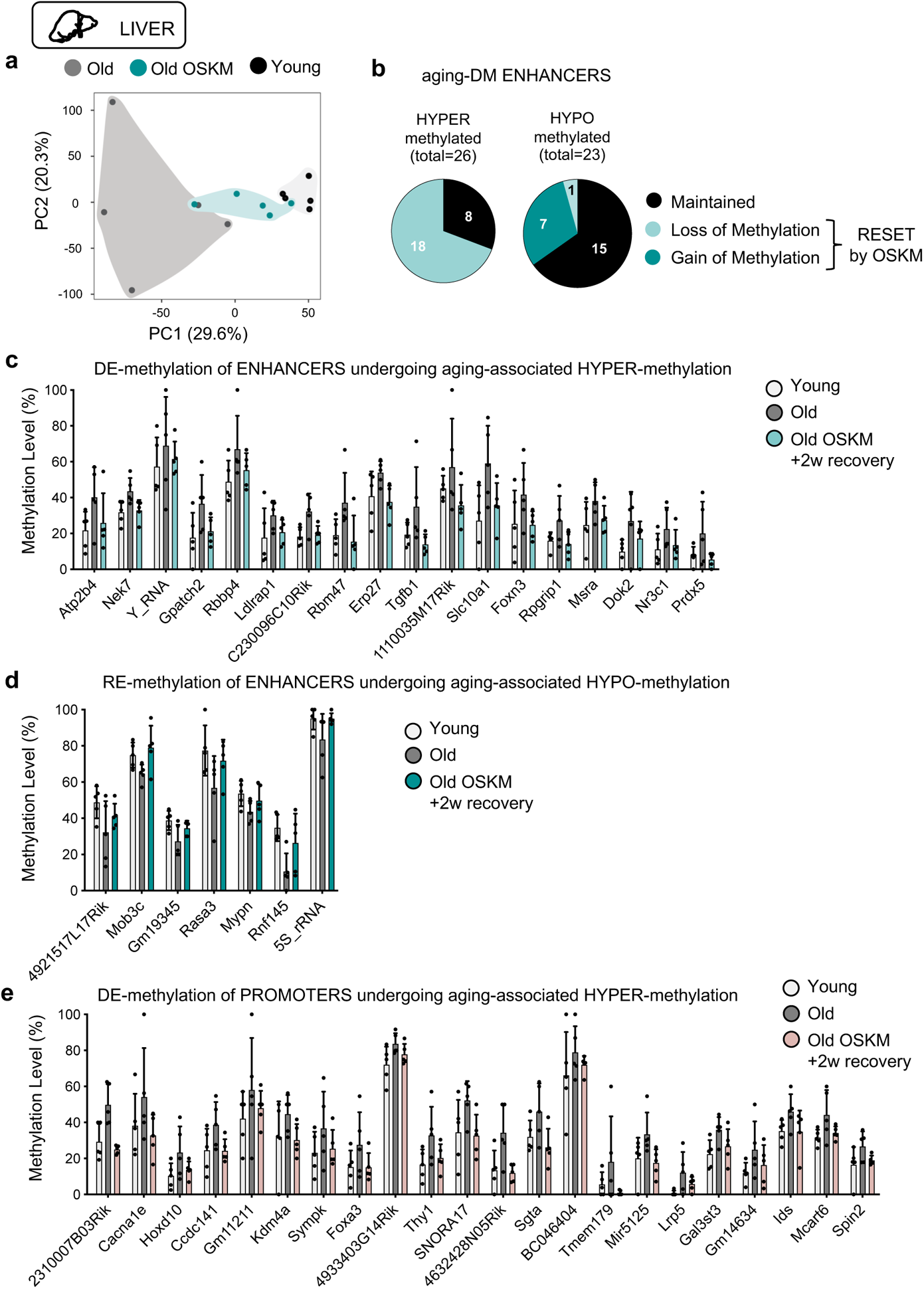
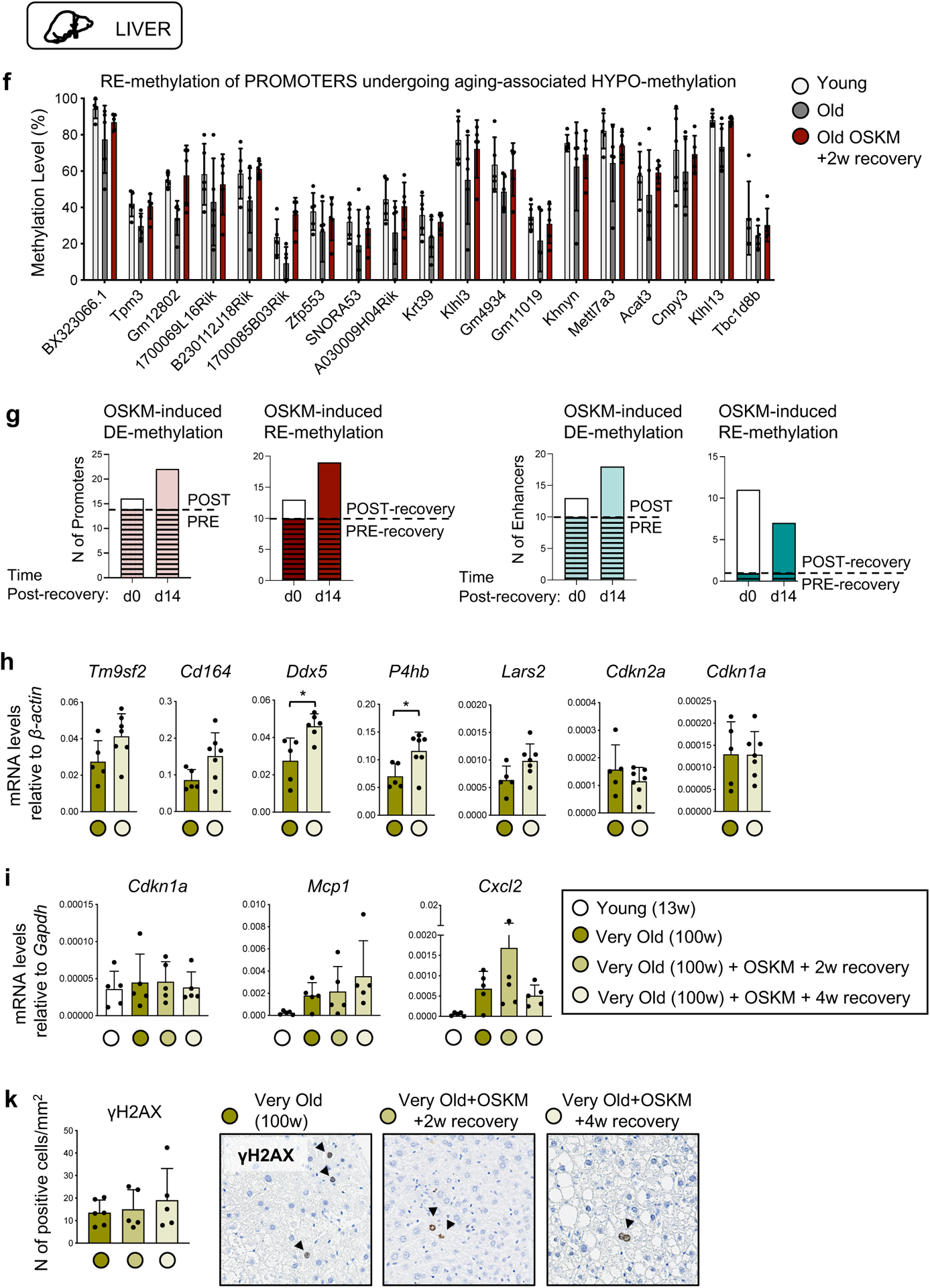
Old livers present rejuvenated features after transient OSKM reprogramming. **a,** PCA of aging-associated DM enhancers in young, old and old-OSKM livers. Old mice are 55 weeks of age. **b,** DM enhancers are classified into hyper- and hypomethylated during aging, and shown is the subset of these enhancers that alters their methylation profile due to transient OSKM activation. **c,** A group of gene enhancers with gain of methylation undergoing aging-associated hypomethylation. **d,** A set of gene enhancers with loss of methylation levels undergoing aging-associated hyper-methylation. **e,** A set of gene promoters with decreased methylation levels after transient OSKM activation undergoing aging-associated hypermethylation in liver. **f,** A group of gene promoters with gain of methylation undergoing aging-associated hypomethylation in liver. **g,** The methylation status of aging-hypermethylated or hypomethylated promoters and enhancers that were found above to be OSKM-demethylated or remethylated respectively was evaluated directly after OSKM cessation (day 0 post-recovery) and 14 days post-recovery in the liver samples. **h,** The expression of global aging genes identified by mouse Aging Cell Atlas^37^, as well as p16 (*Cdkn2a)* and p21 (*Cdkn1a)* expression was evaluated in very old livers (100 weeks; group 2 consists of 5 wild-type mice as control and 7 reprogrammable mice activating OSKM for 1 week and 4 weeks of recovery). **i,** p21 (*Cdkn1a*), *Mcp1* and *Cxcl2* expression in the liver of young (13 weeks) and very old (100 weeks; group 1) mice. k, Immunohistochemistry of γH2AX in the liver of very old (100 weeks) mice. Statistical significance was evaluated using one-way ANOVA with Tukey’s multiple comparison method, and comparisons are indicated as *P < 0.05, **P < 0.01 and ***P < 0.001. Bars in c-f and h-k represent the standard deviation (SD) of the data.

**Extended Data Figure 4.**
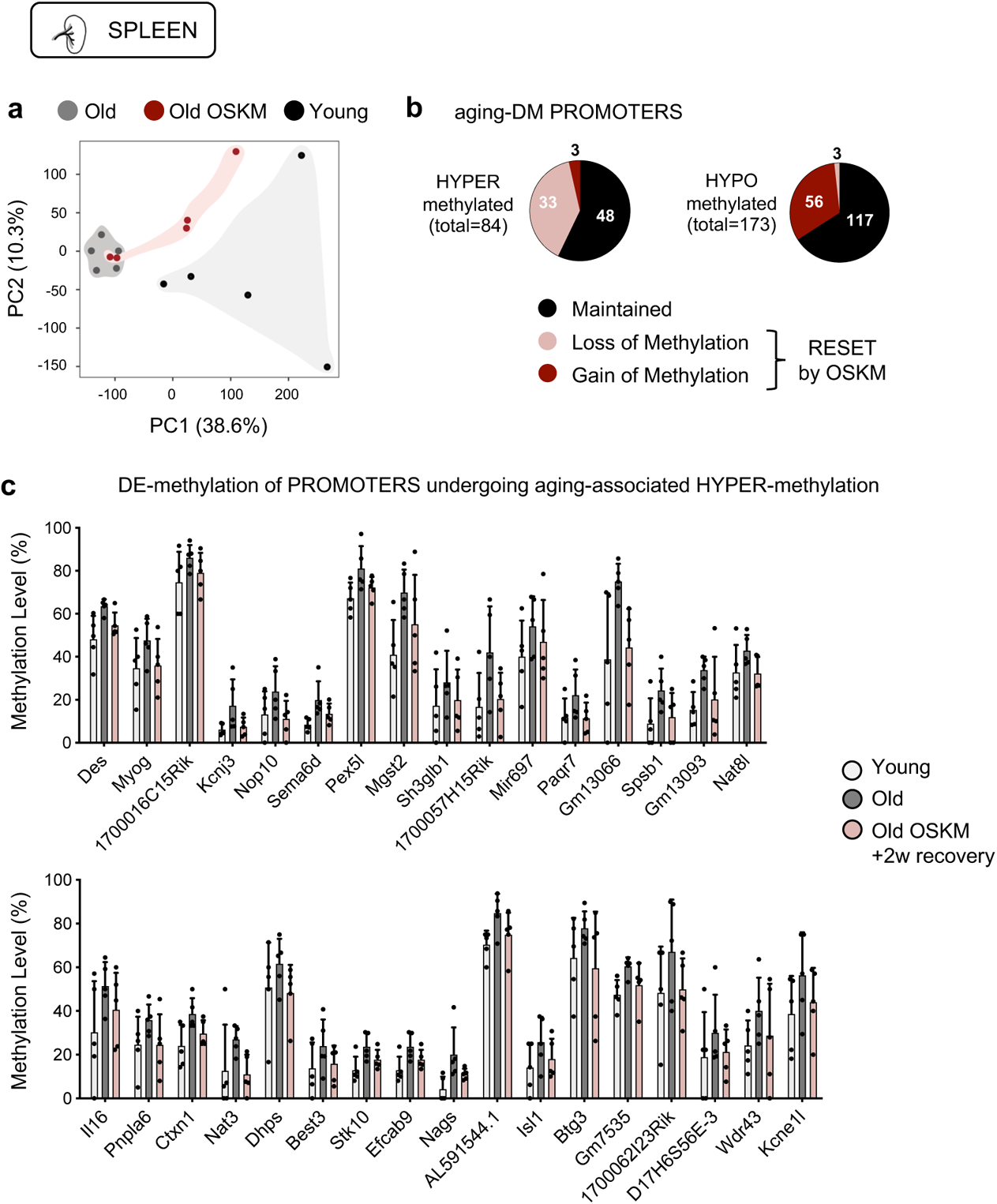
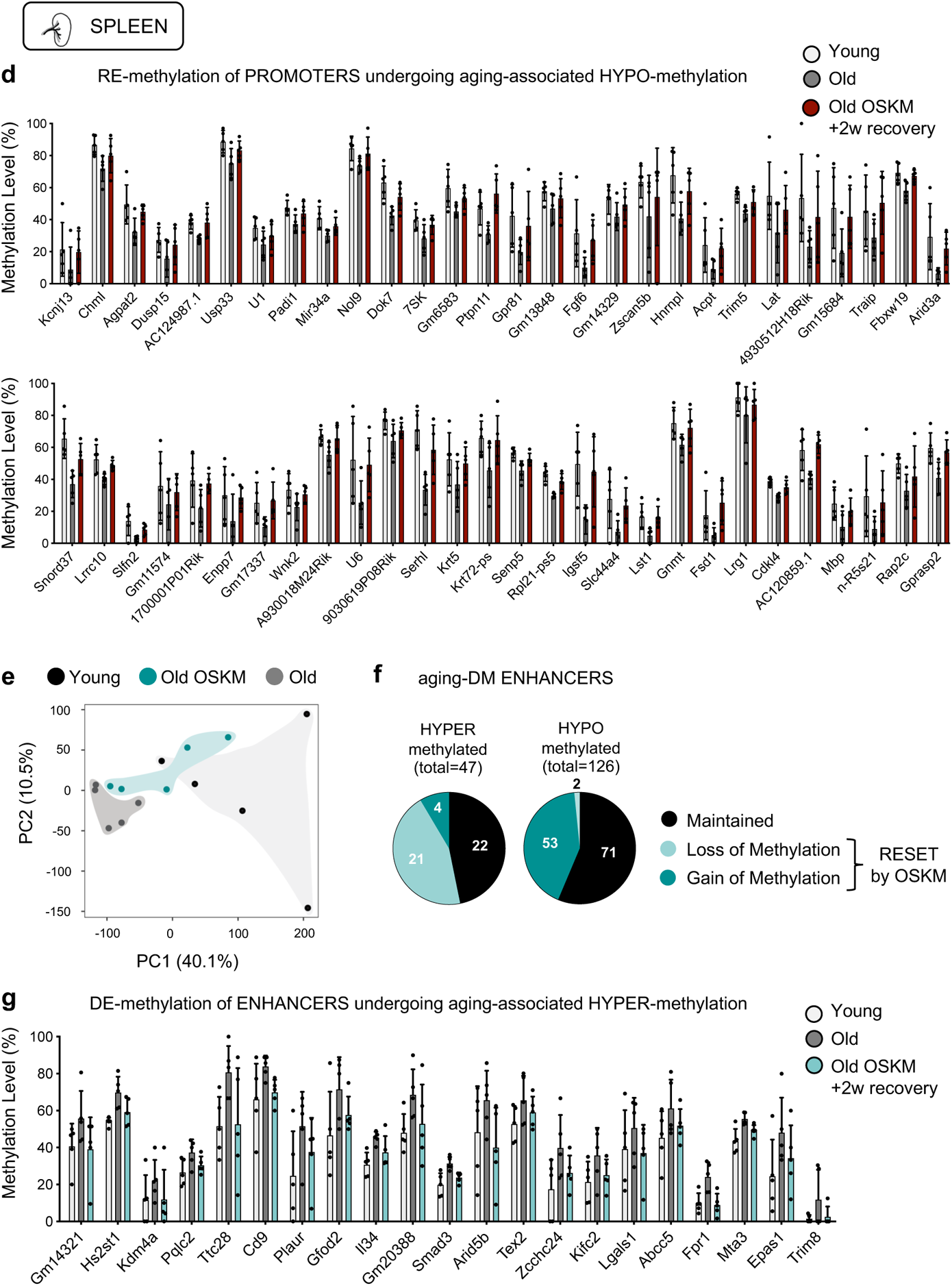
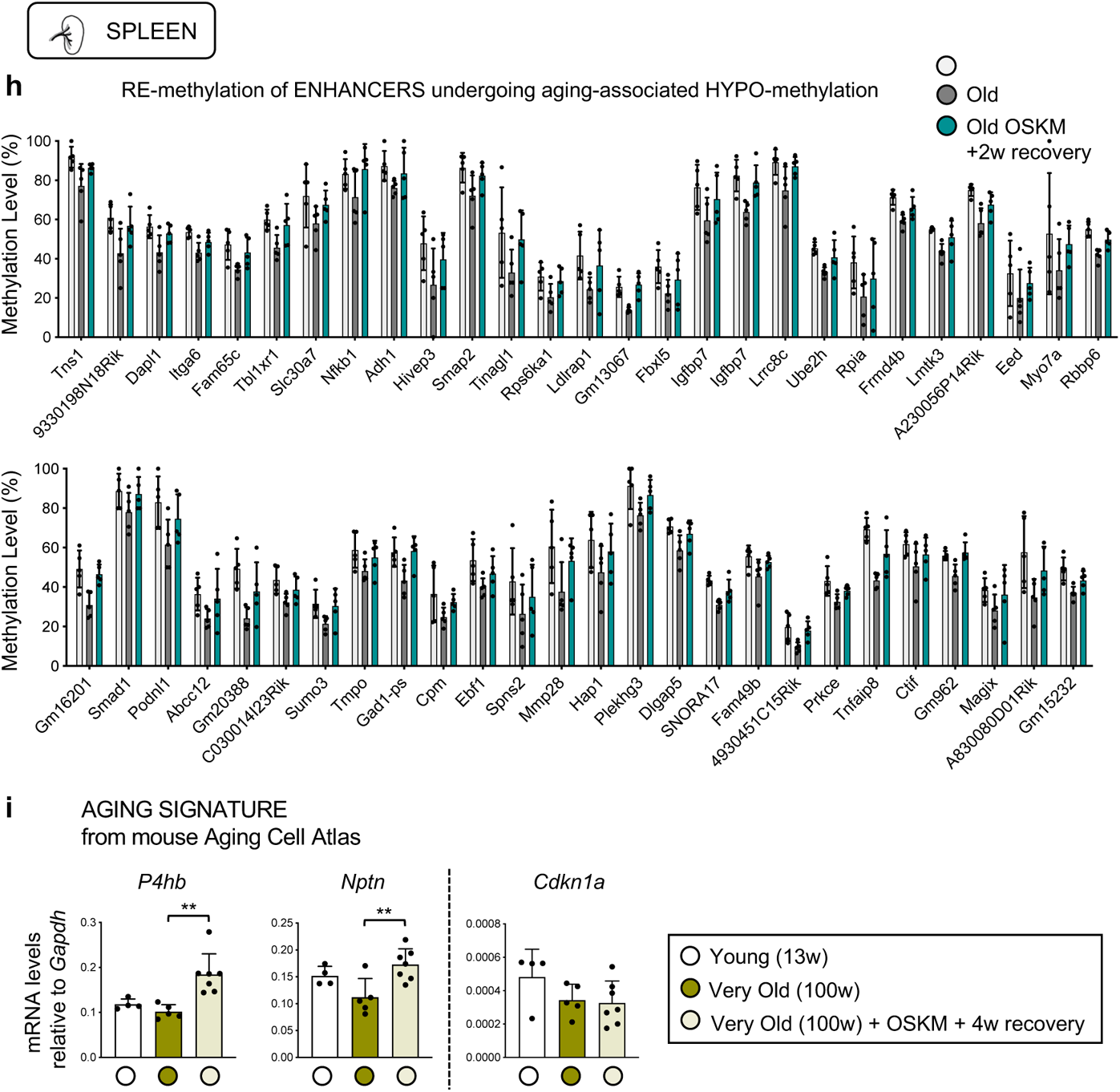
Evidences of OSKM-induced rejuvenation in haemopoietic cells. **a,** Principal Component Analysis (PCA) of aging-related differentially methylated (DM) promoters of young, old and old-OSKM spleens. **b,** DM promoters are classified into hyper- and hypo-methylated during aging, and shown is the number of these promoters that alter their methylation profile due to transient OSKM activation. **c,** A set of gene promoters with decreased methylation levels after transient OSKM activation undergoing aging-associated hypermethylation in spleen. **d,** A group of gene promoters with gain of methylation undergoing aging-associated hypomethylation in spleen. **e,** PCA of aging-associated DM enhancers of young, old and old-OSKM spleen. **f,** DM enhancers are classified into hyper- and hypo-methylated during aging, and shown is the subset of these enhancers that alters their methylation profile due to transient OSKM activation. **g,** A group of gene enhancers with loss of methylation undergoing aging-associated hypermethylation. **h,** A set of gene enhancers with gain of methylation undergoing aging-associated hypomethylation. **i,** The expression of global aging genes identified by mouse Aging Cell Atlas^37^, as well as p21 (*Cdkn1a)* expression was evaluated in very old spleens (100 weeks; 5 wild-type mice as control, 7 reprogrammable mice activating OSKM for 1 week and recovering for 4 weeks) compared to young (13 weeks; n= 4) control spleens. Bars in c-d and g-i represent the standard deviation (SD) of the data.

**Extended Data Figure 5.**
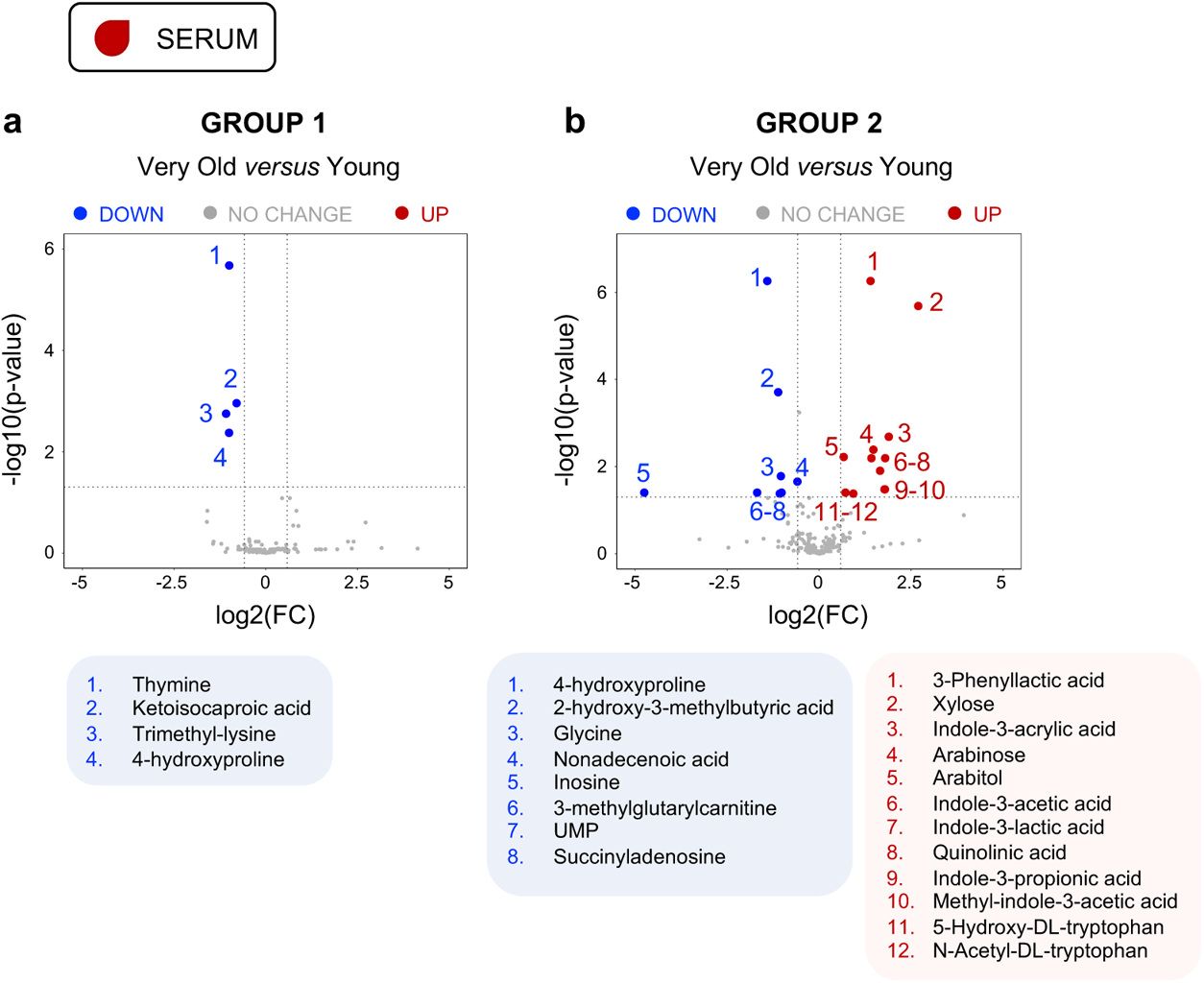
Metabolomic analysis in the serum of very old-OSKM mice. **a,** Volcano plot depicting the differentially present metabolites (fold change>1.5 and adjusted pval<0.05) in the serum of very old (100 weeks, n=6) *versus* young (13 weeks, n=3) female mice as *Group 1*, and **b,** as *Group 2* consisting of very old mice (100 weeks, n=6) versus young (15 weeks, n=5) female mice. Independent metabolomic analyses have been performed for the two different cohorts of mice. Statistical significance was evaluated using a non-parametric linear mixed-effect model. Comparisons are indicated as *P < 0.05, **P < 0.01 and ***P < 0.001.

